# Fmp40 ampylase regulates cell survival upon oxidative stress by controlling Prx1 and Trx3 oxidation

**DOI:** 10.1101/2024.04.20.590396

**Authors:** Suchismita Masanta, Aneta Wiesyk, Chiranjit Panja, Sylwia Pilch, Jaroslaw Ciesla, Marta Sipko, Abhipshita De, Tuguldur Enkhbaatar, Roman Maslanka, Adrianna Skoneczna, Roza Kucharczyk

**Affiliations:** Institute of Biochemistry and Biophysics PAS, Warsaw 02-106, Pawinskiego 5A; Institute of Biology, College of Natural Sciences, University of Rzeszow, Rzeszow, Poland

**Keywords:** Fmp40 ampylase, mitochondrial redoxins, Prx1, Trx3, programmed cell death, yeast

## Abstract

Reactive oxygen species (ROS), play important roles in cellular signaling, nonetheless are toxic at higher concentrations. Cells have many interconnected, overlapped or backup systems to neutralize ROS, but their regulatory mechanisms remain poorly understood. Here, we reveal an essential role for mitochondrial AMPylase Fmp40 from budding yeast in regulating the redox states of mitochondrial 1-Cys peroxiredoxin, Prx1, which is the only protein shown to neutralize H_2_O_2_ with the oxidation of the mitochondrial glutathione and Trx3, thioredoxin, directly involved in the reduction of Prx1. Deletion of *FMP40* impacts a cellular response to H_2_O_2_ treatment that leads to programmed cell death (PCD) induction and an adaptive response involving up or down regulation of genes encoding, among others the catalase Cta1, PCD inducing factor Aif1, and mitochondrial redoxins Trx3 and Grx2. This ultimately perturbs the reduced glutathione and NADPH cellular pools. We further demonstrated that Fmp40 AMPylates Prx1, Trx3, and Grx2 *in vitro* and interacts with Trx3 *in vivo*. AMPylation of the threonine residue 66 in Trx3 is essential for this protein’s proper endogenous level of and its precursor forms’ maturation under oxidative stress conditions. Additionally, we showed the Grx2 involvement in the reduction of Trx3 *in vivo*. Taken together, Fmp40, through control of the reduction of mitochondrial redoxins, regulates the hydrogen peroxide, GSH and NADPH signaling influencing the programmed cell death execution.

**Graphical abstract:** 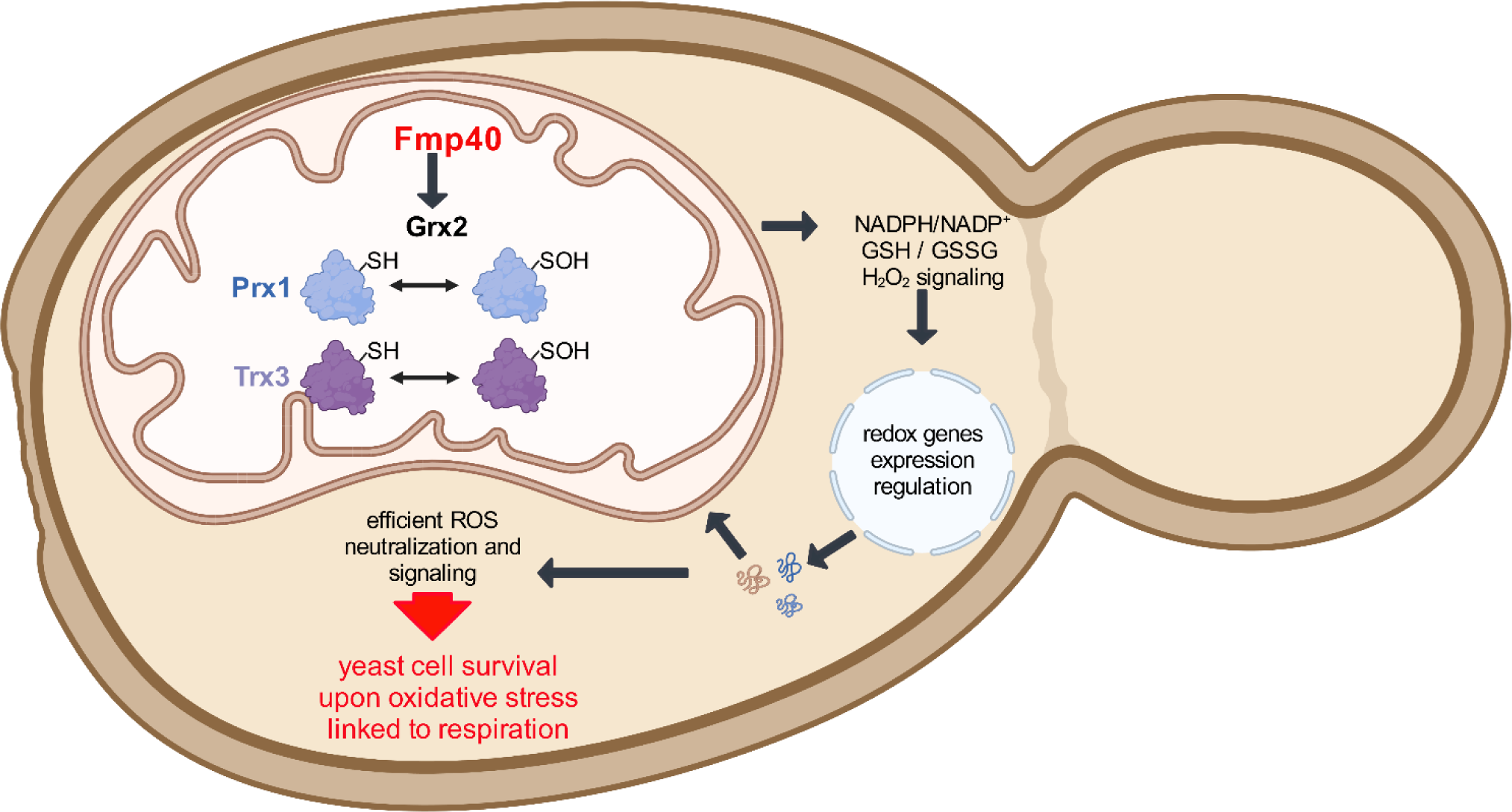

## Introduction

Many reactions taking place in the cell, in particular oxidative phosphorylation in mitochondria or the activity of NADPH oxidases, are accompanied by the production of reactive oxygen species (ROS) ^1–3^. The level of ROS is tightly controlled for two reasons: (i) they are essential signaling molecules regulating crucial cellular processes, among them the induction of programmed cell death (PCD); (ii) they are highly reactive and potentially toxic to the cells’ components, leading to cellular dysfunction and cell death ^4–8^. The mitochondrial inter-membrane space (IMS) in particular, has a high oxidative environment, which defines the course of processes in this mitochondrial sub-compartment ^9^. Several IMS proteins are regulated by oxidation or reduction. Recently we have reported that the yeast IMS-localized Fmp40 protein, belonging to the highly evolutionarily conserved SelO family, is regulated by its oxidation and reduction ^10^. SelO proteins have an AMPylase activity, in which the adenosine monophosphate (AMP) moiety is attached to the threonine, serine, or tyrosine residues in proteins, and are active when they are in a reduced state ^11,12^. Fmp40 is the only identified AMPylase in *Saccharomyces cerevisiae* to date. Yeast cells lacking Fmp40 do not show any growth defects under normal conditions, but their survival upon temporary exposition to oxidative stress triggered by hydrogen peroxide or menadione is decreased ^10^. In *Escherichia coli*, the glutaredoxin GrxA was found to be a substrate of the YdiU SelO AMPylase and the protein S-gluathionylation levels were decreased in bacterial cells as well as in yeast mitochondria ^10^. Therefore, SelO proteins are important regulators of redox enzymes in the cell by AMPylation. However, the substrates and the mechanism of their action are unknown.

Cells have many ROS-neutralizing proteins/systems keeping cellular redox balance and involved in ROS signaling ^13^. Among them are superoxide dismutases or catalases that directly detoxify ROS, while the glutathione (GSH) and thioredoxin systems are involved in this process indirectly ^14^. Glutathione is a tripeptide containing the one thiol group in the cysteine residue, which may be attached to other protein thiols to protect them against irreversible oxidation or may itself become oxidized during reductive reactions while providing the reducing force with the formation of glutathione disulfide (GSSG) ^15^. The GSH system includes the associated enzymes: glutathione reductase, which is necessary to maintain the proper pool of reduced GSH, and enzymes that use GSH as a reductive power for their activity like GSH peroxidases, glutaredoxins, or GSH transferases. Independent from the GSH system are thioredoxins with their reductases using NADPH-regenerating system for thioredoxins reduction ^16^. Both the systems partially overlap as GSH is involved in the reduction of thioredoxins as well ^17,18^. The matrix GSSG level, dependent on the activity of mitochondrial peroxidase Prx1, is an important indicator of matrix redox homeostasis ^19^. The third group of proteins acting as antioxidants or molecular chaperones is peroxiredoxins ^20^.

Mitochondria, like the cytosol, have a complete GSH and thioredoxin/peroxiredoxin systems. The cytosolic glutathione reductase Glr1, catalase Cta1, and superoxide dismutase Sod1 (present in the mitochondrial IMS possess their mitochondrial echoforms, but a superoxide dismutase Sod2 localizes uniquely in the mitochondrial matrix ^21–23^. Yeast contains two cytosolic glutaredoxins, Grx1 and Grx2; however, Grx2 is also present in mitochondria ^24^. Among three yeast monothiol glutaredoxins (Grx3-5), the Grx5 is mitochondrially localized and is required for the maturation of enzymes containing iron-sulfur clusters, e.g., contributing to the believed essential mitochondrial function Fe-S cluster biosynthesis ^25^. The mitochondrial thioredoxin system is made up of Trx3 thioredoxin and Trr2 thioredoxin reductase ^26,27^. Among three yeast glutathione peroxidases Gpx1, Gpx2, and Gpx3 the last two have mitochondrial echoforms ^28^. In the *in vitro* experiments, the purified Grx1 and Grx2 also have glutathione peroxidase activity and can reduce hydroperoxides directly ^29,30^. Additionally, mitochondria possess a peroxiredoxin Prx1, crucial in maintaining mitochondrial matrix redox balance ^31,32^. Prx1 is the only protein for which the efficient reaction with H_2_O_2_ has been proven ^19^.

Prx1 is a single 1-cysteine peroxiredoxin, which has a dual mitochondrial localization: in the IMS, where the mature protein starts from the 31st amino acid residue, and in the mitochondrial matrix, where the mature protein is deprived of the first 38 amino acid residues resulting in the one cysteine peroxidase ^33^. Respiration or exposure to hydrogen peroxide enhances Prx1 expression ^34^. The Prx1 reduction requires the cooperation of different antioxidant systems, including mitochondrial thioredoxin Trx3, Trr2, and GSH ^19,35–37^. The peroxidatic Cys_91_ residue of Prx1 is detected in a reduced form unless cells are subjected to oxidative stress. Hydroperoxides oxidize Cys_91_ to the sulfenic acid form, which can then be glutathionylated through a reaction with GSH ^36^. This mixed disulfide is a substrate for reduction by Trr2, in a reaction that proceeds through the formation of a Prx1-Trr2 disulfide-bonded intermediate, regenerating reduced active Prx1. The GSH-sulfenic acid form of Prx1 can be reduced by the formation of a Prx1-Trx3 disulfide-bonded intermediate as well ^38^. An additional role in Prx1 reduction has been suggested for mitochondrial glutaredoxin Grx2, based on the observation that Grx2 can reduce glutathionylated Prx1 *in vitro* but was not demonstrated *in vivo* ^35^.

Here, we investigated the Fmp40 role in the regulation of the redox state of mitochondrial proteins involved in the neutralization of ROS. We showed that Fmp40 is involved in apoptosis execution, through regulation of the redox cycle of Prx1, the oxidation of which limits the oxidation of matrix GSH to GSSG, an important determinant of cell death. Fmp40 AMPylates *in vitro* mitochondrial redoxins Prx1, Trx3, and Grx2, interacts with Trx3 *in vivo* and impacts the Trx3 redox cycle as well but not the redox states of the Grx2. We also for first time identified the threonine at position 66 in Trx3 is AMPylated. Modification of this residue is important for the regulation of Trx3 protein levels and maturation of the Trx3 precursor protein under oxidative stress. Additionally, we confirmed that Grx2 is involved in the reduction of the thioredoxin Trx3 *in vivo*, regardless of the presence or absence of Fmp40, postulating that the primary function of Grx2 in Prx1 reduction is fulfilled at the level of Trx3 reduction.

## 2. Materials and methods

### 2.1. Yeast strains construction

The yeast strains used in the study are listed in Table S1. For genetic interaction analysis, the BY4741 strain background was used. Strains from yeast knock-out collection were obtained from Open Biosystems. The double mutants with the *FMP40* gene deletion were constructed by crossing the respective single deletion mutants in *MATa* background with *MATα fmp40Δ::kanMX4*, sporulation of the diploid cells, and tetrads dissection. Deletion of a given gene in the MR6 background was introduced to the yeast genome by transformation with a specific deletion cassette, amplified with *gene*-Up and *gene*-Low primers (Table S2) using the total DNA from the respective yeast knock-out strain, selection of transformants on the geneticin containing plates and PCR verification. The same approach was applied to the construction of the double mutants in the MR6 strain background. MR6 strain is a derivative of the W303-1B strain, in which the nuclear *ARG8* gene was replaced with *HIS3*, containing the mitochondrial genome of the S288C strain that has been entirely sequenced, and has a wild-type *CAN1* gene, encoding a basic amino acid permease ^39^. The strain expressing the Trx3-3xMyc tagged variant was kindly provided by Prof. Chris Grant (University of Manchester, UK, ^26^). The *TRX3-3MYC* gene variant was introduced into *fmp40Δ, prx1Δ, grx2Δ*, *prx1Δ fmp40Δ*, and g*rx2Δ fmp40Δ* strains in the MR6 background by crossing these strains with the isogenic but of the opposite mating type *TRX3-3MYC* strain, sporulation of obtained diploids and tetrads dissection. The correctness of all genetic constructs was validated by PCR using appropriate primers and Verif-KanMx primer (Table S2).

For the introduction of point mutations into the *TRX3-T_66_* codon in the yeast genome, the plasmids pSM32 and pSM33 (see plasmids description below), containing *URA3* marker and *TRX3* sequences encoding the Trx3-T_66_>A and Trx3-T_66_>E proteins, respectively were cut with the MscI restriction enzyme within the *TRX3* sequence. The linearized plasmids were introduced into the wild-type MR6 and BY4741 strains. The uracil prototrophic colonies were then passaged on the 5-FOA medium to select cells that removed the *URA3* gene. The presence of the mutation in *TRX3* gene was verified by sequencing of PCR amplified *TRX3* gene sequence with Trx3-For and Trx3-Low primers (Table S2). To introduce the HA tag to Trx3, Trx3-T_66_>A, and Trx3-T_66_>E, the PCR-based strategy was applied ^40^. The cassette obtained using Trx3-3HA-Up and Trx3-3HA-Low primers (Table S2) and the pFA6a-3HA-kanMX6 plasmid as a template ^41^ was introduced into MR6, RKY358, and RKY359 strains’ by high-efficiency transformation. Similarly, the Cta1 and Ctt1 were labeled with a 3HA tag. The integration cassettes were amplified with primers Ctt1lower, Ctt1upper, or Cta1lower, Cta1upper (Table S2). The reaction products were introduced into the MR6 strain by high-efficiency transformation. Clones with gene fusions were selected on YPGluA+geneticin plates. The correctness of clones was proved by Western Blot using the anti-HA antibody (clone 16b12, Babco Inc).

### 2.2. Growth media and conditions

Media for growing yeast were: rich glucose YPGlu/YPGluA (1% Bacto yeast extract, 1% Bacto Peptone, 2% glucose, additionally, 40 mg/L adenine in YPGluA), rich glycerol YPGly/YPGlyA (1% Bacto yeast extract, 1% Bacto Peptone, 2% glycerol, 40 mg/L adenine in YPGlyA), SC selective medium: (6.7 g/L YNB w/o amino acids, 2% glucose or glycerol, 40 mg/L adenine, appropriate amino-acids drop-out powder. Media were solidified by the addition of 2% (w/v) Bacto agar. Media powders of Difco^TM^ were purchased from Thermo Fisher Scientific, amino-acids drop-out powders from Formedium. Yeast cells were grown at 28 or 36 °C, and cultures in liquid media were shaken at 180 rpm. Growth curves were established with the Bioscreen CTM system. Geneticin G418 (Thermo Fisher Scientific) was added to the YPGluA plates at the concentration of 200 µg/mL. Hydrogen peroxide (Sigma-Aldrich) was added to the solid media cooled to 50 °C at the concentration indicated in the figure legends.

For colony forming units (CFU) assay yeast cells were cultured in a liquid YPGluA, YPGlyA, or SC selective media at 28 °C until OD_600_=2. The culture was divided into two parts of equal volume. The H_2_O_2_ in appropriate concentrations depending on the medium (YPGlyA - 0.03 mM H_2_O_2_; YPGluA - 0.2 mM H_2_O_2_; W0 - 0.1 mM H_2_O_2_; determined experimentally, given by the survival of the control strain at the level of ∼ 50-80%) was added to one of the samples. Both cultures were incubated for 200 min with shaking. After incubation, a series of ten-fold dilutions of both cultures were prepared and plated on a solid medium containing glucose as a carbon source (to plate approximately 100 cells per dish). After 2 days of incubation at 28 °C, the colonies that had grown on the plates were counted and the percentage of cells that survived the H_2_O_2_ treatment was calculated. The number of colonies that grew in the control conditions allowed us to calculate the initial number of cells, taken as 100%. The experiment was performed in three biological repetitions.

For qPCR, steady state, and redox states of proteins analysis, cells (pre-grown in YPGluA liquid medium) were grown in YPGlyA medium to OD_600_=2-3. The cultures were divided into three parts of equal volume. The H_2_O_2_ was added at two concentrations to two cultures (0.03 and 0.3 mM), and then all cultures were incubated for 200 minutes with shaking and processed for protein or RNA extraction.

For the drop-test growth analysis, cells were pre-grown in the YPGluA liquid medium, serially diluted, and then 5 µL of cells’ suspension was spotted on the plates. Plates were incubated at two temperatures 28 or 36 °C. Plates were photographed daily from the second to the fifth day of incubation.

### 2.3. Protein purification, in vitro AMPylation assays, and generation of the antibodies

The Prx1, Trx3, Tsa2, and Grx2 proteins were produced using pET15b-based plasmids encoding these proteins (kindly provided by Prof. Antonio Barcena) and following the protocols described in the references ^35,42,43^. The purification of Fmp40 protein and *in vitro* AMPylation reactions were performed according to the protocols described in detail in reference ^10^. Briefly, Prx1, Trx3, Grx2 or Tsa2 purified protein (4 µg) was incubated with Fmp40 (1 µg) in the AMPylation buffer (50 mM Tris-HCl pH 8,5 mM MgCl_2_, 5 mM DTT, 100 µM [α−^32^P]ATP (SA = 500–3000 cpm/pmol)) for 30 minutes at 37 °C and terminated by addition of EDTA to 25 mM concentration. After the addition of SDS loading buffer to the samples, they were boiled. Reaction products were resolved using SDS-PAGE. Gel was stained with Coomassie and air dried overnight before exposition to a high-sensitive X-ray film or a radioactivity-storage screen (Fujifilm). The film was developed after one week depending on the intensity of the radioactive signal, the signal from the screen was visualized using a Phosphor imager (FujiFilm FLA 5100). For non-radioactive AMPylation reactions, 100 μM ATP was used in the reactions. After EDTA termination and the methanol-chloroform precipitation, protein modifications were analyzed by mass spectrometry. The Fmp40, Prx1, and Grx2 specific antibodies were generated using purified proteins from *E. coli* extracts as antigens. The immunization of rabbits, isolation, and affinity purification of polyclonal antibodies were performed by Davids Biotechnologie GmbH (Germany).

### 2.4. Site-directed mutagenesis of Prx1 and Trx3

The Q5 Site-Directed Mutagenesis Kit (New England Biolabs) was used to introduce the mutations into pET15b plasmid-borne copies of *PRX1* and *TRX3* genes using pairs of adequate, specific primers: Prx1-S116A-Up, Prx1-S116A-Low and Trx3-T95A-Up, Trx3-T95A-Low, according to the manufacturer protocol. A similar protocol was applied to introduce a mutation into the 66th codon of the *TRX3* gene. The coding sequence of *TRX3* was subcloned from the pRS416-TRX3 plasmid (received from Prof. C. Grant) into the pRS306 plasmid ^44^ with KpnI-SacI restriction enzymes, giving the plasmid pSM31. The sequences of primers used for mutagenesis, namely, Trx3-Mut-left, Trx3-T66E, or Trx3-T66A are listed in Table S2. The resulting plasmids, pSM32 and pSM33, encode the Trx3-T66>A and Trx3-T66>E proteins, respectively.

### 2.5. RNA extraction, cDNA synthesis, and quantitative real-time PCR

For total RNA extraction, the Bead-Beat Total RNA Mini kit (A&A Biotechnology) was used. The RNA was extracted from 8 x 10^7^ yeast cells and homogenized using a TissueLyser LT bead mill (Qiagen). Residual DNA was removed by digestion with TURBO™ DNase (Invitrogen) according to the manufacturer’s instructions. The 1.8 µg of total RNA was reversed-transcribed into cDNA in a 20 µL reaction mixture, using RevertedAid H Minus First Strand cDNA Synthesis Kit (Thermo Fisher Scientific) according to the manufacturer’s instructions. Obtained cDNA was precipitated by adding 1/10 volume of 3 M sodium acetate, pH 5.2, and 2.5 volumes of 96% ethanol, washed 85%, 80%, and 75% ethanol, and dissolved in 10 µL of water. The cDNA concentration was estimated using a NanoDrop ND-1000 spectrophotometer (Thermo Fisher Scientific). The real-time quantitative PCR assays were carried out on the PikoReal 96 Real-Time PCR System (Thermo Fisher Scientific). Primers were designed using the Primer Quest program (http://eu.idtdna.com/PrimerQuest/Home/Index). The reaction mixture contained: 1x concentrated commercial SYBR Green mixture 2xHS-PCR Master Mix SYBR A (A&A Biotechnology), forward and reverse specific primers (100 nM of each), cDNA template (in three concentrations, each in technical duplicates), and water to 10 µL of the final volume. Reactions were performed with an initial denaturation step of 95 °C for 1 minute, followed by 50 cycles of denaturation (95 °C for 12 seconds) and primer annealing extension (68 °C for 30 seconds). Fluorescence was read during the annealing-extension step of each cycle. After cycling, melting point temperature analysis was performed in the range of 60 °C to 95 °C with temperature increments of 0.2 °C. The quality of results was evaluated based on expected Ct differences among three cDNA amounts as well as product melting curves. Rare outlying results were omitted from calculations. Three concentrations of cDNA allowed us to calculate individual efficiencies for each primer pair and normalize all results to one, common for all genes, cDNA concentration. The amount of each target gene was calculated by the modified ΔCt method with a geometric mean of two reference genes Cts as reference ^45^. We used the Grubbs test (alpha = 0.05) to remove outliers. The Welch’s t-student test was applied to check whether the observed differences between strain and conditions were statistically significant.

### 2.6. Co-immunoprecipitation of mitochondrial proteins

The mitochondria were prepared from yeast cells grown in a liquid-rich glycerol medium at 28 °C, by the enzymatic method described in reference ^46^. The 1000 µg of mitochondria was centrifuged at 9500 x g for 10 minutes and suspended in 400 µL of the solubilization buffer (20 mM Tris/HCl pH 7.4, 0.1 mM EDTA, 50 mM NaCl, 10% [vol/vol] glycerol, 1% [wt/vol] digitonin, 2 mM PMSF, 1 x EDTA free proteinase inhibitor (Roche, Indianapolis, IN)) and incubated for 30 minutes on ice. For Trx3-Myc protein purification and subsequent mass spectrometry analysis, after centrifugation at 9500 x g for 10 minutes, the supernatant was collected and incubated with 200 µL of Pierce Anti-c-Myc Magnetic beads (Thermo Fisher Scientific). For the co-immunoprecipitation experiment, a DSP cross-linker (Sigma-Aldrich) was added to the final concentration of 2 mM, and the supernatant was further incubated for 30 minutes at 25 °C. The cross-linker was quenched by the addition of Tris pH 7.4 to the concentration of 25 mM for 15 minutes. The 60 µL of the supernatant was taken for the SDS-PAGE analysis while 340 µL of the supernatant was incubated with 50 µL of Pierce Anti-c-Myc Magnetic beads (Thermo Fisher Scientific) at 4 °C, on a rotator. The bound proteins were analyzed by mass spectrometry or eluted from beads by boiling them in 50 µL of Laemmli sample buffer for 5 minutes. The 20 µL of the mitochondrial extract (input) and 20 µL of bead eluates were loaded per lane on SDS-PAGE gels, 12%, and 16%.

### 2.7. Protein extraction and Western blotting

For SDS-PAGE analysis 10 OD of cells were centrifuged and suspended in 500 µL of 0.2 M NaOH. After 10 min incubation on ice, the samples were mixed with 50 µL of 50% TCA (trichloroacetic acid), incubated 10 min on ice, and centrifuged at 21950 x g for 10 min at 4 °C. The protein pellet was washed with 1 mL of 1 M Tris-base and suspended in 50 µL of 5% SDS, and the concentration of proteins was measured by the method of Lowry ^47^. The 50 µg of proteins were loaded per lane and run on 10 to 18% gels, depending on the protein size ^48^. The proteins were then transferred onto a nitrocellulose membrane using iBlot2 Gel Transfer Device (Thermo Fisher Scientific) or a semi-dry protein transfer system from BIO-RAD and analyzed by Western blotting. Briefly, the membrane was blocked with 5% BSA or 5% milk for 1 hour at room temperature, rinsed with PBST (50 mM Tris-HCl pH 7.5, 150 mM NaCl, 0.1% Tween 20), and incubated with indicated in the figure legends primary antibodies. The anti-Pgk1 (Thermo Fisher Scientific) or anti-Por1 polyclonal antibodies (a kind gift from Prof. Teresa Żołądek, IBB PAS) were used for normalization. For immunodetection, the incubation with a respective secondary antibody conjugated to horseradish peroxidase (1:10000 in PBST, DAKO) for 1 hour at room temperature was performed, followed by 3–4 washes with PBST. The blots were then sealed with Immobilon Western Chemiluminescent HRP Substrate from Millipore, used as a substrate for HRP.

### 2.8. Determination of the oxidation state of Trx3 and Prx1 in vivo

The redox state of Trx3 was measured by covalent modification with the thiol-reactive probe 4-acetamido-4’maleimidyldystilbene-2,2’-disulfonic acid (AMS; Molecular Probes) as described previously ^49^. The binding of AMS to one cysteine adds about 0.5 kDa to the protein mass, which yields a shift in the migration compared to the oxidized protein. The addition of 1 kDa to 14.4 kDa Trx3 protein, having two cysteines, is enough to make a visible shift on 18% SDS-PAGE gels. To perform an assay, the 10 OD of cells were collected by centrifugation, and suspended in 200 μl of 10% TCA. Sterile glass beads of 0.45 mm diameter (equal to the volume of the cell pellet) were added to the cells’ suspension, and cells were homogenized three times for 30 seconds, using a Mini-BeadBeater (Biospec Product, USA). The homogenates were then centrifuged for 10 min at 9500 x g and the pellets were washed with 1 M Tris-HCl, pH 8.8. The resulting pellets were suspended in 50 µL of 2x non-reducing Laemmli sample buffer containing Complete Protease Inhibitors and 10 mM PMSF with or without 25 mM AMS or in the reducing sample buffer containing 25 mM AMS, and 10 mM TCEP. Samples were incubated for 15 min in an ice bath, then 10 minutes at 37 °C, and finally 2 minutes at 100 °C. Because mature Prx1 of 29.5 kDa contains only one cysteine, to distinguish the redox states of this cysteine, a mmPEG_24_-based alkylation was applied, yielding a shift of ∼ 2.4 kDa in the migration of reduced Prx1 ^19,50^. Briefly, yeast cells were grown to the mid-log phase, and the 10 OD of cells were harvested by centrifugation at 1500 × g for 3 minutes at room temperature and washed with isosmotic buffer. Pellets were suspended in 100 µL SDS-loading buffer containing either 10 mM mmPEG_24_ (steady state), DMSO (unmodified), or 10 mM mmPEG_24_ and 10 mM TCEP (maximally reduced) and boiled for 5 min at 100 °C. The cells were then disrupted as above in the dark. After an additional 60 minutes incubation in the dark 8 µL of samples were analyzed by Western blotting against Prx1.

### 2.9. Annexin V apoptosis assay

To measure the apoptosis we used an annexin V protocol adapted for yeast cells by Kaushal et al. ^51^. Briefly, the *fmp40Δ* and wild-type strains in BY4741 background were grown at 28 °C in YPGlu medium with shaking to an early exponential growth phase (∼5 x 10^6^ cells / mL), divided into two parts, from which the first served as control. In the second part, the H_2_O_2_ was added to the final concentration of 2.5 mM to induce apoptosis. After three hours of incubation in the same conditions, the 10^8^ cells from each culture were collected, centrifuged, rinsed with water, and digested with Zymolyase 100T in buffer S (1.2 M sorbitol, 0.5 mM MgCl_2_, 35 mM KPi, pH 6.5), until when spheroplasts were ready. After washing with binding buffer (0.8 M sorbitol, 10 mM HEPES/NaOH, pH 7.4, 140 mM NaCl, 2.5 mM CaCl_2_, 0.1% BSA), equal amounts of the spheroplasts’ suspension were taken for staining using Annexin V Apoptosis Detection Kit FITC (Invitrogen). The first stain was performed using Annexin V conjugated to FITC for 15 minutes in the dark. Then, after twice washing with the same buffer, propidium iodide was added to the samples. Analysis was performed using FACSCalibur (Becton Dickinson) using the FL1 for Annexin V labeled with FITC and FL2 channel for propidium iodide. Three independent biological repetitions were made for both tested strains, with 30000 cells analyzed per sample.

### 2.10. Fluorescence microscopy

The BiG Mito-Split-GFP strain was transformed with pAG414pGPDβ11 expressing Fmp40-β11, Pgk1-β11 or Atp4-β11 [2]. Cells were grown in SC-glycerol-ura liquid medium till OD_600_=2. For tracking co-localization of Nuc1 with nucleus marker, the pNUC1-GFP (kindly provided by dr Aneta Kaniak-Golik) and pWJ1323 [NUP49-CFP] plasmids were used ^52^. For tracking co-localization of Nuc1 with mitochondrial marker, the pNUC1-GFP and pYX142-mtRFPff plasmids were used, respectively ^53^. Strains of the wild-type and *fmp40Δ* in BY4741 background carrying chosen plasmids were grown in SC glucose medium lacking uracil with shaking at 28 °C to an early exponential phase (∼5 x 10^6^ cells / mL). Then, two 2 mL samples per culture were taken, centrifuged (1000 x g), and suspended in 2 mL of the same fresh medium (control) or the medium supplemented with H_2_O_2_ to 2.5 mM final concentration (pro-apoptotic conditions). Samples were then incubated for 200 min at 28 °C in a rotating wheel. After incubation, 1 mL of each sample was centrifuged, and the pellet was suspended in 50 µL of PBS. The 2 µL of cells’ suspension was then placed on microscope slides and analyzed using Zeiss Axio Imager M2 microscope, operated by Zeiss Axio Vision 4.8 software, under 100 x magnification (using Zeiss EC Plan-NeoFluar/1.3 NA objective lens) under immersion oil. Imaging was performed using a Zeiss AxioCam MRc5 Digital Camera (Zeiss, Oberkochen, Germany). The Nuc1-GFP, Nup49-CFP, and mtRFP signals were visualized using the Zeiss 38HE, 49HE, and 63HE filter sets respectively and DIC (for bright field). Three biological repetitions were performed for every strain and condition, number of cells analyzed in each condition is given in the legend of the corresponding figure. Fluorescent microscopy images quantitative analysis was performed using ImageJ Fiji software, and Cell Counter and DeconvolutionLab2 plugins, with the Richardson-Lucy algorithm (15 iterations) on fluorescent channels to correct a systematic error of the optical system used in the experiment. A number of cells was counted based on DIC channel images; the fluorescent signals were counted in all used in the certain experiment channels. After the counting was done fluorescent channels were merged into a single image and a number of cells with overlapping fluorescent proteins signals were counted. Data obtained that way was analyzed in RStudio software. To see whether variables are dependent or not on each other, data was divided into four groups representing wild-type control cells, wild-type cells after H_2_O_2_ treatment, *fmp40Δ* control cells, *fmp40Δ* cells after H_2_O_2_ treatment. Each group was associated with a categorical variable defined either by the absence or presence of co-localization of Nuc1-GFP and Nup49-CFP (or Nuc1-GFP and mtRFP). A contingency table was created for each pair of variables, and the χ^2^-dependency test was carried out.

### 2.11. Determination of glutathione content

The glutathione content (total, GSH, and GSSG levels) was determined in the yeast cells with GSH/GSSG-Glo Assay according to the manufacturer’s protocol (Promega) adapted for yeast cells as was done by Kwolek-Mirek et al. ^54^. Briefly, yeast cells of analyzed strains were cultured in a liquid YPGlyA medium at 28 °C until OD_600_=1-2. The culture was divided into two parts of equal volume. The H_2_O_2_ at 0.3 mM concentration was added to one culture, then both cultures were incubated for 200 minutes with shaking. After incubation, cell density (number of cells per mL) was determined using the Malassez chamber. A cell number of 5 × 10^5^ suspended in PBS buffer was used for assay. From each culture, cell suspension was added to a white flat bottom 96-well plate in duplicate, one for measuring total glutathione (GSH total) and the second for measuring oxidized glutathione (GSSG) level. All samples were treated for 10 minutes with intensity shaking with the manufacturer’s Lysis Reagent containing Luciferin-NT. The reduced glutathione (GSH) in samples used for GSSG determination was blocked by NEM (N-Ethylomaleimide). After lysis, all samples in a 1:1 ratio were incubated for 30 minutes with Luciferin Generation Reagent containing glutathione-S-transferase. After incubation, 100 µL of Luciferin Detection Reagent containing stabilized luciferase was added to all samples. The luminescence signal was proportional to the amount of formed luciferin and thus the amount of GSH was recorded after 15 min using an Infinite 200 microplate reader (Tecan Group Ltd., Männedorf, Switzerland). The total glutathione and GSSG concentrations were read based on standard curves, whereas the level of GSH was calculated by subtracting the GSSG from the total glutathione concentration (due to 1 mole of GSSG is generated by 2 moles of GSH the values of GSSG were multiplied by 2).

### 2.12. Determination of NADP(H) Content

NADP(H) content in the yeast cells was assessed with NADP/NADPH-Glo Assay according to the manufacturer’s protocol (Promega) with the modification described by Kwolek-Mirek et al. ^54^. Briefly, yeast cells were cultured in a liquid YPGlyA medium at 28 °C until OD_600_=1-2. The culture was divided into two parts of equal volume. The H_2_O_2_ at 0.3 mM concentration was added to one culture, then both cultures were incubated for 200 minutes with shaking. After incubation, cell density (number of cells per mL) was determined using the Malassez chamber. The density of 2 × 10^6^ cells/mL in PBS was used for assay. From each culture, the cell suspension was transferred to Eppendorf tubes and incubated in a 1:1 ratio with lysis solution (0.2 M NaOH with 1% DTAB (dodecyl trimethylammonium bromide) for 15 minutes with intense shaking. After incubation, samples were split into separate tubes for acid (measuring NADP^+^) and base treatments (measuring NADPH). To measure NADP^+^ samples were treated with 0.4 M HCl in ratio 2:1; samples for NADPH were without additional compounds. Both types of samples were incubated at 60 °C for 15 minutes. After incubation samples were cooled down to room temperature and neutralized to the same buffer formulation (addition of 0.5 M Tris to acid-treated samples or addition of 0.5 M Tris/HCl to base-treated samples). After preparation 100 µL of both types of samples (acid or base-treated) was added to a white flat bottom 96-well plate and mixed with 100 µL of freshly prepared NADP/NADPH-Glo Detection Reagent containing proluciferin and luciferase. The luminescence signal proportional to the amount of NADP^+^ or NADPH was recorded for 2 hours at 25 °C using an Infinite 200 microplate reader (Tecan Group Ltd., Männedorf, Switzerland). The rate of luminescence increase per minute was measured based on the straight part of luminescence kinetic. The value of the blank was subtracted each time. The results were presented as individual (NADPH and NADP^+^) cofactors content and the calculated NADPH/NADP^+^ ratio.

### 2.13. Statistical analysis

All the experiments were performed in at least three biological repetitions. The statistical tests used for each experiment are specified in the appropriate method section or the figure legend.

### 2.14. Data analysis

When not specified in the figure legends, for image acquisition the Uvitec Cambridge Q4 Alliance System (Uvitec Cambridge, UK) and the manufacturer’s software were used. The images were collected in the serial accumulation option. Images were processed with Adobe Photoshop CC 2019. All figures in this study were assembled using Microsoft Office PowerPoint or Adobe Photoshop (Adobe Inc.). For quantification of pixels in the bands of proteins the ImageJ software was used (www.imagej.net/ij). The raw MS data were pre-processed with Mascot Distiller (version 2.8, Matrix Science) and proteins were identified with Mascot Server (version 2.8, Matrix Science) using the SGD database (6,713 sequences; 3,019,540 residues). The parameters were as follows - enzyme: semiTrypsin, instrument: HCD, missed cleavages: 2, fixed modification - Methylthio (C), variable modifications: Oxidation (M), Phospho (STY), Phosphoadenosine (KSTY). Each file was re-calibrated offline, resulting typical peptide mass tolerance of 5 ppm and fragment mass tolerance of 0.01 Da. FDR was calculated using a target-decoy strategy, threshold of 5% was used.

## 3. Results

### 3.1. Fmp40 promotes apoptosis

The published data about SelO proteins from different organisms suggested that these proteins regulate redox homeostasis and signaling ^55–57^. Thus, we asked about *fmp40Δ* strain sensitivity to oxidative stress induced by growing them under graded concentrations of hydrogen peroxide. The test revealed that the growth of the *fmp40Δ* strain did not differ from the wild-type strain up to 3 mM H_2_O_2_ concentration in the YPGlu medium; however, at higher concentrations, the *fmp40Δ* cells showed better survival than wild-type cells (Fig. 1A). Since the 3 mM H_2_O_2_ is, in fact, a pro-apoptotic hydrogen peroxide dose ^58,59^, we asked if the presence of *FMP40* may influence the PCD.

**Fig. 1.**
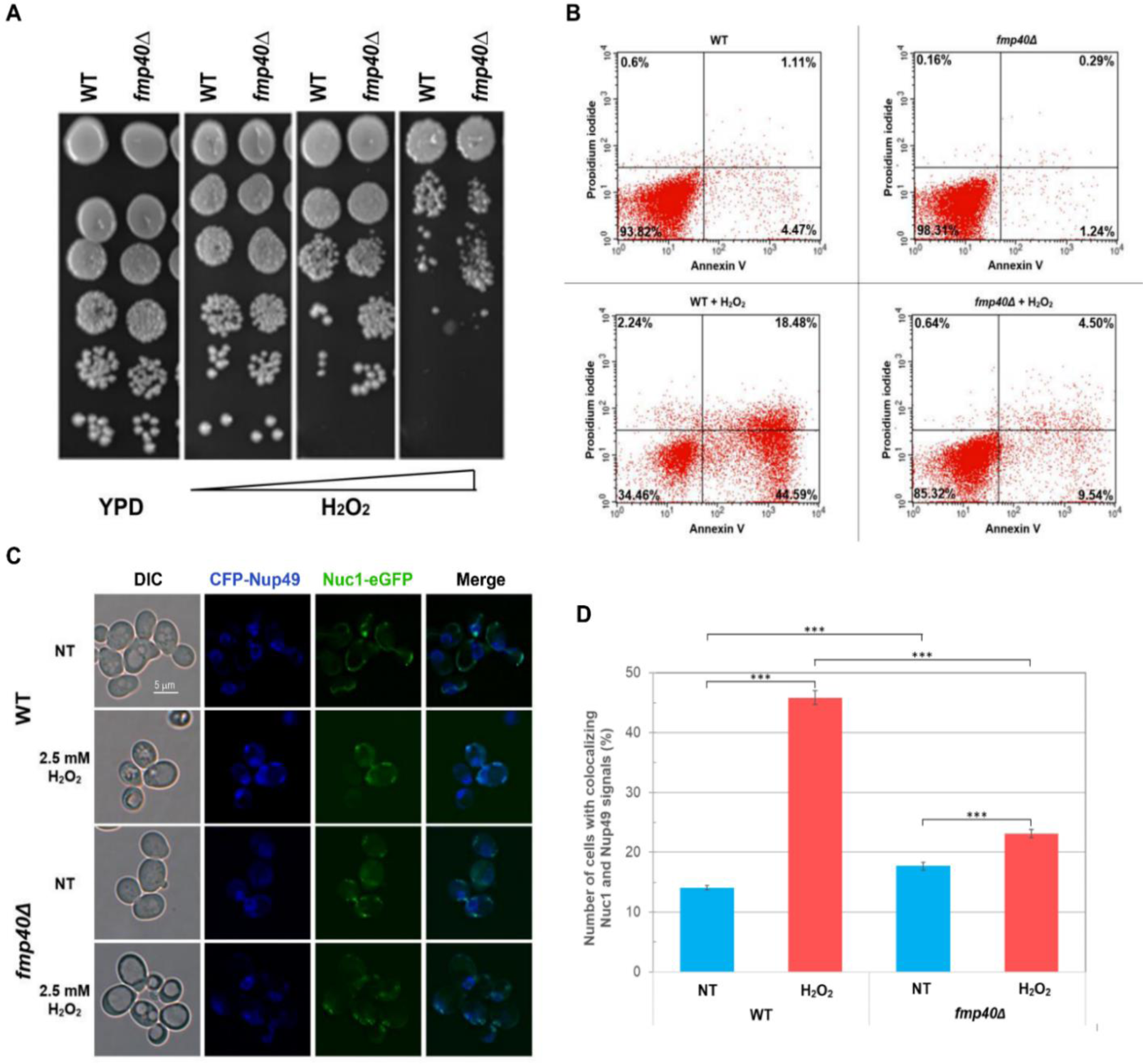
Cells lacking the *FMP40* gene showed a decrease in PCD. **(A)** The growth of *fmp40Δ* cells in the BY4741 background in the presence of H_2_O_2_. Serially diluted cells were spotted on the YPGlu plates containing growing concentrations of H_2_O_2_ (3, 4, and 5 mM, respectively). Growth of the cells was documented after three days of incubation at 28 °C. **(B)** Annexin V assay. Apoptosis in exponentially growing cells (upper panel) of WT (BY4741) (left panel) and *fmp40Δ* (right panel) strains was induced by incubation with 2.5 mM H_2_O_2_ per 200 min in YPGlu medium with shaking (see Material and Methods section for details). Representative results of flow cytometry experiment using propidium iodide as necrotic cells indicator and annexin V as apoptotic cells indicator were shown. **(C)** The same strains as in (A & B) but carrying plasmids pWJ1323 [*CFP::NUP49*] ^52^ and pNuc1-GFP [*TEF1pNUC1::eGFP*] were cultivated to exponential growth phase in minimal selection medium (SC-Ura-His) with shaking, then treated with 2.5 mM H_2_O_2_ to induce apoptosis. Examples of cellular localization of Nuc1-eGFP and nucleus marker CFP-Nup49, characteristic for tested strains and conditions, were acquired using fluorescence microscopy. **(D)** The frequency of Nuc1 and Nup49 colocalization was quantified based on fluorescent microscopy studies. Approximately 1800 cells per strain and condition were analyzed. The cells showing both fluorescent signals were considered only. NT – non-treated cells. To check if the results were statistically significant, the χ2-dependency test was applied using RStudio software and R programming language. P-values (p-value < 0.001 in all pairs of variables) indicate that there is a statistically significant association between variables.

To verify the PCD markers, firstly, we performed an annexin V/propidium iodide (PI) staining assay. This method allows the measurement of a number of apoptotic and necrotic cells in the given population after pro-apoptotic treatment (e.g., hydrogen peroxide treatment). With the addition of annexin V to control cells, almost no fluorescent signal arises. The exposition of phosphatidylserine, which accompanies the PCD process, allows its binding with annexin V linked to the fluorescent dye, generating a fluorescent signal detectable by flow cytometry. Staining cells with PI in parallel enables visualization of necrotic cells present in the assayed population (Fig. 1 B). The percentage of apoptotic cells (the right panels of each graph) was approximately 5.5% (+/- 1.5) for the WT, 1.5% (+/- 0.6) for *fmp40Δ* cells in the control conditions and grew to 63% (+/- 9) and 14% (+/- 5), respectively, after PCD induction with 200 minutes treatment with 2.5 mM H_2_O_2_.

One of the phenotypes accompanying PCD is a Nuc1 nuclease relocation from mitochondria to the nucleus ^60^. Recently, the role of Nuc1 nuclease as the enzyme directly involved in triggering PCD was questioned ^51^, but translocation of Nuc1 from mitochondria to the nucleus under pro-apoptotic conditions is so a characteristic phenomenon for this kind of stress that we decided to employ the test in our research. Hence, we asked whether the translocation of Nuc1 to the nucleus would be changed after treatment of cells lacking *FMP40* with pro-apoptotic H_2_O_2_ concentration. Results shown in Fig. 1 C, D documented a decreased number of cells with Nuc1-eGFP and nucleus marker CFP-Nup49 co-localization, after PCD induction with hydrogen peroxide compared to wild-type control. However, the initial number of cells showing co-localization of traced proteins in control conditions was higher in the *fmp40Δ* cells population. Thus Fmp40 protein is needed for PCD induction.

It is known that the morphology of the mitochondrial network changes in various stress conditions ^61^. We noticed that the Nuc1-eGFP signal looked different in oxidative stress than in control conditions, and those differences were not limited to protein translocation to the nucleus. To discriminate the localization of the green fluorescent signal we observed, we used not only Nuc1-eGFP but also two fluorescent markers, CFP-Nup49 and mtRFPff. For this goal, we transformed wild-type (BY4741) and *fmp40Δ* strains with three plasmids, pNuc1-GFP, pWJ1323 [*CFP::NUP49*] ^52^, and pYX142-mt-RFPff ^53^ and checked co-localization of fluorescent organelles’ markers and Nuc1-GFP. The results of this experiment presented in Fig. S1 confirmed a dual, mitochondrial, and nuclear localization of Nuc1-eGFP, as well as the fact that nuclease Nuc1 more frequently co-localized with CFP-Nup49, e.g., was present in the nucleus, in the wild-type cells subjected to oxidative stress than if stress was applied to the cells lacking *FMP40*. The experiment also allowed associating the Nuc1-eGFP signals with their mitochondrial localization, even when the mitochondrial network underwent structural changes due to their fission upon oxidative stress conditions.

### 3.2. Genetic interactions of fmp40Δ with genes encoding proteins of cellular redox systems

Genetic interactions have long been studied in various model organisms to identify functional relationships among genes or their corresponding products. Because SelO family proteins were suggested to influence the redox homeostasis, and we showed that one of ROS–dependent phenotype, e.g., PCD induction is changed in yeast cells lacking Fmp40, we decided to take a closer look at the epistatic relation between *FMP40* and genes encoding proteins involved in defense against ROS, mainly but not exclusively focusing on mitochondrial proteins. Based on information from literature and those deposited in the SGD database, we selected genes encoding proteins engaged in cellular redox systems or protection against oxidative damage and removed the *FMP40* gene from the strains lacking chosen genes (Table S1). Due to the availability of yeast knock-out collection ^62^ and the hydrogen peroxide resistance phenotype of *fmp40Δ* cells in the drop-test (Fig. 1A) we conducted genetic tests in the BY4741 strain background. The drop sensitivity test was performed on a YPGlu solid medium supplemented with rising concentrations of hydrogen peroxide (3, 4, and 5 mM, respectively) at 28°C (Supplementary Fig. S2). Considering the number of colonies growing on the spots, where serially diluted cells’ suspensions were dropped on plates, we gathered information concerning the survival of cells treated with hydrogen peroxide. The sensitivity test revealed that some of the analyzed mutants were more sensitive or resistant to applied oxidative stress than the wild-type control strain. The strains’ sensitivity to oxidative stress varied in strength and depended on the stressor concentration. The rate of sensitivity versus wild-type was displayed on the graph in Fig. 2. The single *yap1Δ, gpx3Δ, grx2Δ, tsa2Δ, sod2Δ, srx1Δ, glr1Δ, grx5Δ, aif1Δ, tsa1Δ, sod1Δ, ctt1Δ, cta1Δ, prx1Δ, pos5Δ*, and *ycp4Δ* strains showed sensitivity, while *ahp1Δ, oxr1Δ, dot5Δ, mgp12Δ, grx3Δ, aim6Δ, fmp46Δ, gpx2Δ, gsh1Δ, mdl2Δ, gtt2Δ, oye2Δ, gsh2Δ, trx1Δ, trr2Δ, gpx1Δ, aim33Δ*, and *aim14Δ* strains showed resistance to applied concentrations of H_2_O_2_ in comparison to the control strain. Removal of the *FMP40* gene from analyzed mutants allowed to divide them into six groups, according to their phenotypes: (1) *sensitive, Fmp40 independent* (*yap1Δ, gpx3Δ, grx2Δ, tsa2Δ*); (2) *sensitive, sensitized by fmp40Δ* (*sod2Δ, srx1Δ, glr1Δ, grx5Δ, aif1Δ, tsa1Δ, sod1Δ, ctt1Δ, cta1Δ*); (3) *sensitive, sensitivity rescued by fmp40Δ* (*prx1Δ, pos5Δ,* and *ycp4Δ*); (4) *resistant, sensitized by fmp40Δ* (*ahp1Δ, oxr1Δ, dot5Δ, mgp12Δ, grx3Δ*), (5) *resistant, resistance increased by fmp40Δ* (*aim6Δ, gsh2Δ* and *fmp46Δ*); (6) *resistant, Fmp40 independent* (*aim14Δ, aim33Δ*, *gpx2Δ, gsh1Δ, gtt2Δ, mdl2Δ, oye2Δ, trx1Δ, trr2Δ*, and *gpx1Δ*). The *trx3Δ* mutant behaved like the wild-type control strain. The most pronounced effect of H_2_O_2_ on the strain viability was visible for *yap1Δ* from group 1, *sod2Δ, grx5Δ, srx1Δ, glr1Δ* from group 2, and *pos5Δ* from group 3. The most evident dependence on *fmp40Δ* showed mutants from groups 3 and 4 since *FMP40* gene deletion in their background led to a reversed phenotype than observed in a single mutant treated with H_2_O_2_. Some strains showed delayed growth in the tested conditions, as the size of the colonies formed by these strains was smaller than those of the control strain. Such phenotype was observed for *tsa1Δ*, *grx5Δ* already in the control conditions, and for *pos5Δ*, *ycp4Δ*, *grx2Δ*, *prx1Δ*, *glr1Δ*, *sod1Δ*, *srx1Δ* on medium with hydrogen peroxide. These complex interactions between *FMP40* and the above genes indicate a primary function of Fmp40 in the regulation of cellular systems that control the balance of NADP/NADPH, GSH/GSSG, Fe-S clusters biosynthesis, and neutralize reactive oxygen species, in particular the hydrogen peroxide.

**Fig. 2.**
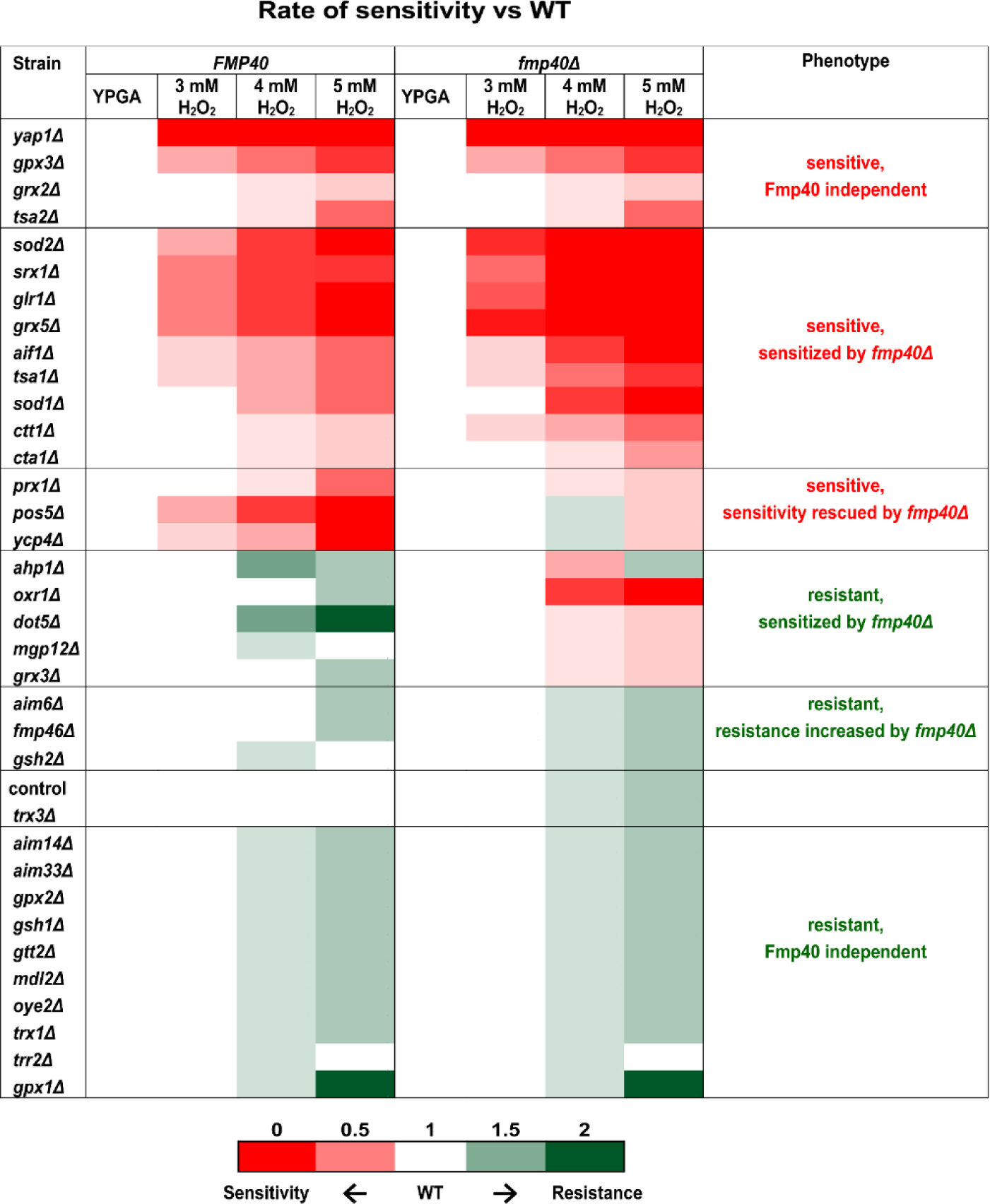
The genetic interactions of *fmp40Δ* with genes encoding known redox proteins assessed by the growth phenotype in the presence of hydrogen peroxide at 28 °C - classification of genes according to the growth phenotype in relation to the presence of *FMP40* gene. The pictures of the plates are shown in supplementary Fig. S2.

### 3.3. Changes in mRNA levels of genes encoding proteins that neutralize reactive oxygen species in fmp40Δ cells

The hydrogen peroxide resistance phenotype of *fmp40Δ* cells in the BY4741 background was surprising, because previously, in the W303-1B background, we observed a decrease in the survival of *fmp40Δ* cells after exposure to lower concentrations of hydrogen peroxide – 0.2 mM or 0.03 mM in YPGluA or YPGlyA media, respectively, while in the drop test we did not observe any differences in growth (^10^, Fig. S3). To understand this phenotypic difference, which may be linked to the genetic background, we checked the expression level of forty genes encoding main components of the glutathione and redox defense systems (including genes for which we observed phenotypic changes while testing genetic interactions with *fmp40Δ* in the oxidative stress conditions), as well as catalases, superoxide dismutases and four genes encoding transcription factors known to be involved in controlling the expression of those genes (in W303-1B background ^17^). Because the accumulation levels of Fmp40 increase by at least 10 fold when yeast cells are growing in media containing non-fermentable carbon sources ^10^, we used cells grown in rich glycerol (YPGlyA) medium, exposed to 0.03 or 0.3 mM hydrogen peroxide for 200 minutes, for analysis of mRNA levels of those genes by quantitative qPCR. The 0.03 mM hydrogen peroxide concentration in rich glycerol media led to a decrease in the *fmp40Δ* cells’ survival to a similar extent as the 0.2 mM hydrogen peroxide concentration in the rich glucose medium (Fig. S3B). Therefore, as the high concentration of hydrogen peroxide, in experiments when cells were grown in rich glycerol media, the ten times higher, e.g., 0.3 mM H_2_O_2_ concentration was chosen, by analogy to the growth conditions in rich glucose media, in which the H_2_O_2_ concentration in a range 2-2.5 mM induces the programmed cell death ^59^. The changes in mRNA levels were analyzed comparatively, concerning the tested strains and conditions, which resulted in clusters shown in Fig. 3.

**Fig. 3.**
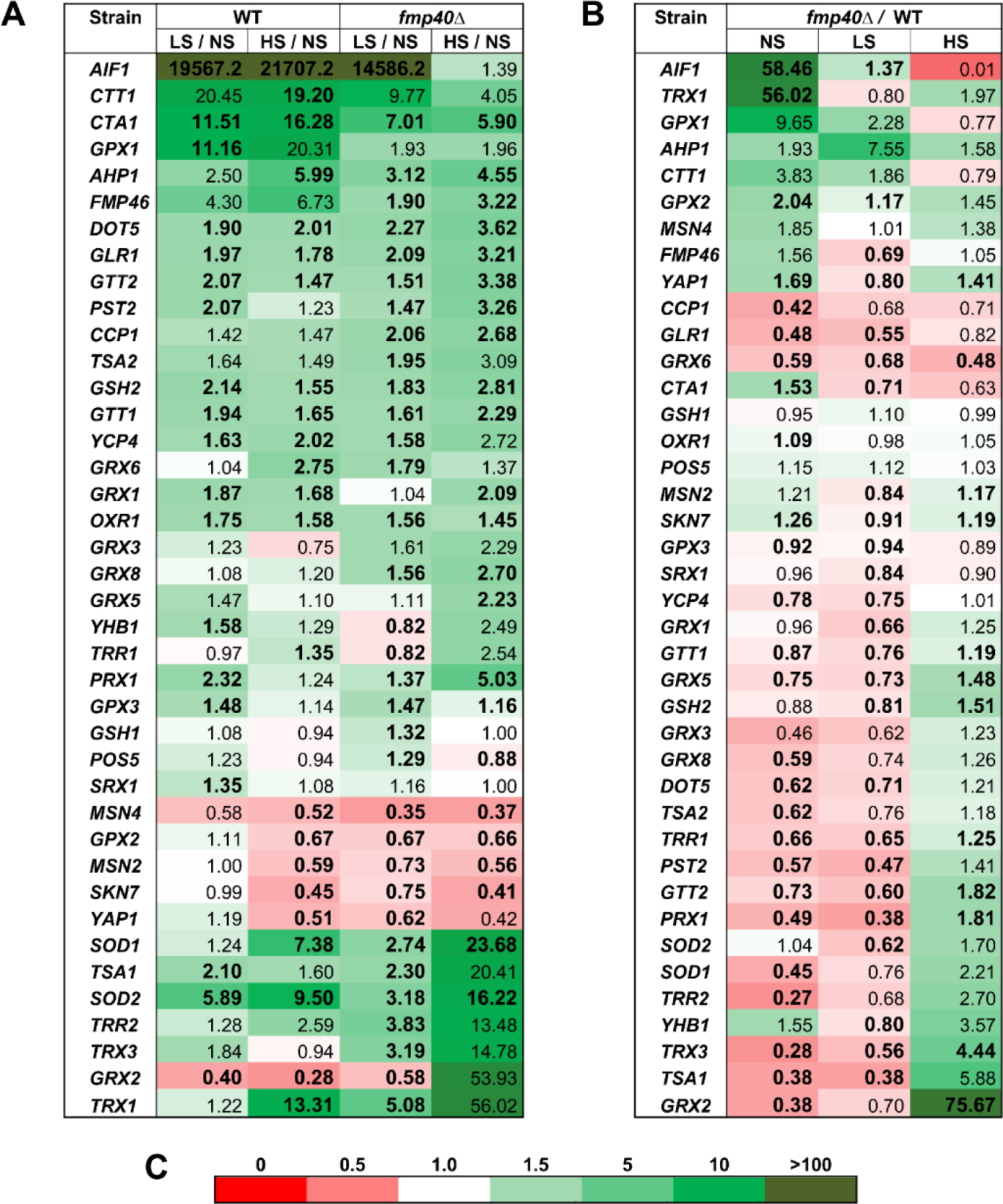
The profile of mRNA expression of the yeast redox or redox-related genes in *fmp40Δ* and control cells depending on stress conditions. The ratio of expression of a given gene calculated with respect to applied conditions (**A**) and strains (**B**) is shown, where (**C**) presents the scale used. The conditions are indicated NS - no stress, LS - 0.03 mM H_2_O_2_, HS - 0.3 mM H_2_O_2_. To visualize the differences in the basal levels of analyzed genes’ expression, the means from at least three biological repetitions for both strains in control conditions were shown. The more the expression of a given gene in the *fmp40Δ* background increased compared to the wild-type, the darker green was used to highlight the ratio. As opposed, the more the expression of a given gene in the *fmp40Δ* background decreased compared to the wild-type, the darker red was used to highlight the ratio (**C**). The results with statistical significance lower than p-value < 0.05 were shown with bigger sizes and bold fonts (Welch’s t-student test was applied). The Euclidean distance method of hierarchical linkage of genes, offered by Gene Cluster 3.0 software, was used for gene clustering.

The experiment has revealed:

1. Changes in genes expression in the control strain upon hydrogen peroxide treatment (Figure 3 A, C) The most pronounced increase in the expression level in response to hydrogen peroxide treatment was noticed for the *AIF1* gene. However, the transcript for this gene is hardly present in non-treated cells therefore the calculated expression rate is artificially elevated. In response to hydrogen peroxide treatment, the expression of *CTA1*, *CTT1*, *GPX1*, and *SOD2* genes increased significantly (several times). We noticed a slight (but more than 1.5 times) increase for *GRX1*, *GTT1*, *OXR1*, *DOT5*, *GSH2*, *GTT2*, *PST2*, *GLR1*, *YCP4*, *PRX1*, *YHB1*, and *TSA1* in response to both concentration of hydrogen peroxide used in the experiment, while for *AHP1*, *GRX6*, and *SOD1* only at higher H_2_O_2_ concentration. In contrast, the expression of a few genes decreased after treatment with the 0.3 mM hydrogen peroxide, and those are genes encoding the transcription factors engaged in the stress response, namely *MSN4*, *YAP1*, *MSN2*, *SKN7*, glutaredoxins *GRX2* and *GRX3*, and glutathione peroxidase *GPX2*.
2. Changes in genes expression in the *fmp40Δ* cells upon hydrogen peroxide treatment (Figure 3 A, C) We noticed several phenotypic categories of analyzed genes regarding their expression changes in response to a growing concentration of hydrogen peroxide in the cells lacking Fmp40. The majority of the analyzed genes showed an increase in expression with increasing hydrogen peroxide concentration; the most substantial for *PRX1*, *SOD1*, *SOD2*, *TSA1*, *TRR2*, *TRX1*, and *TRX3*. Some genes showed a lower increase in the expression level when treated with higher than with a lower concentration of hydrogen peroxide, e.g., *AIF1*, *CTT1*, and *CTA1*. The treatment with low hydrogen peroxide concentration resulted in the reduction of expression of *YHB1*, *TRR1*, and *GRX2*. Finally, *MSN2*, *MSN4*, *GPX2*, *SKN7*, and *YAP1*, all displayed reduced expression when *fmp40Δ* cells were treated with hydrogen peroxide.
3. Changes in genes expression in the *fmp40Δ* cells in comparison to control strain (Figure 3 B, C) From the most significant changes in the expression level of analyzed genes in the *fmp40Δ* background versus wild-type, we detected an increase in expression for *AIF1* and *TRX1* (over 50-times)*, GPX2*, *YAP1* and *CTA1* (more than 1.5 times) and a significant decrease (more than twice) for *TRR2, TRX3*, *TSA1*, *GRX2*, *CCP1*, *SOD1*, *GLR1*, *PRX1* and smaller but statistically significant decrease (less than twice) for *PST2*, *GRX6*, *GRX8*, *DOT5*, *TSA2*, *TRR1*, *GTT2*, *GRX5*, *YCP4*, *GTT1*, and *TRR1*.
4. Changes in genes expression in the *fmp40Δ* strain versus control cells upon hydrogen peroxide treatment (Figure 3 A, C) The expression of the *CCP1*, *GLR1*, and *GRX6* genes decreased in cells lacking *FMP40* gene in control growth conditions and upon both concentrations of hydrogen peroxide treatment. In contrast, expression of *GRX2*, *TRR2, TRX3*, *TSA1*, *SOD1*, *PRX1*, *PST2*, *GRX8*, *DOT5*, *TSA2*, *TRR1*, *GTT2*, *GRX5*, *YCP4*, *GTT1*, *TRR1*, as well as *YHB1* and *SOD2* was decreased in the cells lacking the *FMP40* gene upon treatment with low concentration of hydrogen peroxide and increased upon treatment with higher concentration of hydrogen peroxide. For the *GRX2* gene, the difference in expression was the most pronounced. For some genes, we saw their decreased expression in the *fmp40Δ* background in only one of the tested conditions, e.g., the *CTA1* gene had increased expression compared to wild-type cells in the control conditions, while its expression decreased with exposition of *fmp40Δ* cells on the growing H_2_O_2_ concentration. The expression of the *AIF1*, initially very high, decreased but was still elevated upon the lower concentration of H_2_O_2_; in the higher stress conditions, it was significantly reduced. A similar tendency was observed for *GPX1* and *CTT1* genes. The results of this experiment demonstrated that the control of redox genes expression in the absence of *FMP40* is lost. Levels of mRNA for genes such as *GRX2, TRX1*, and *AIF1* were changed the most significantly in particular conditions. These differential changes in expression patterns of genes encoding redox proteins revealed in the *fmp40Δ* cells explain the opposing susceptibility phenotypes obtained in the viability tests upon exposure to different concentrations of hydrogen peroxide.

### 3.4. Changes in mRNA levels are not always reflected in protein accumulation

After assessing the mRNA levels, we checked the steady-state levels of chosen redox proteins in the same experimental conditions. The accumulation of Prx1 was in accordance with the *PRX1* mRNA levels changes - in non-stressed *fmp40Δ* cells was decreased in comparison to the control cells and after exposition to hydrogen peroxide amount of Prx1 increased similarly in wild-type and *fmp40Δ* cells, independently on the hydrogen peroxide concentration (Fig. 4). In *fmp40Δ* non-stressed cells, the levels of Cta1 and Ctt1 proteins were increased in proportion to the observed increase in the mRNA levels for their genes. Exposition to hydrogen peroxide led to an increase in the levels of both catalases, similarly in wild-type and *fmp40Δ* cells. Thus, the *PRX1, CTT1*, *and CTA1* mRNA changes in non-stressed *fmp40Δ* cells were reflected in the level of the Prx1, Ctt1, or Cta1 proteins, while in cells exposed to hydrogen peroxide were not. Immunodetection with anti-Fmp40 antibody showed that the level of Fmp40 AMPylase increases up to four-fold upon hydrogen peroxide treatment.

**Fig. 4.**
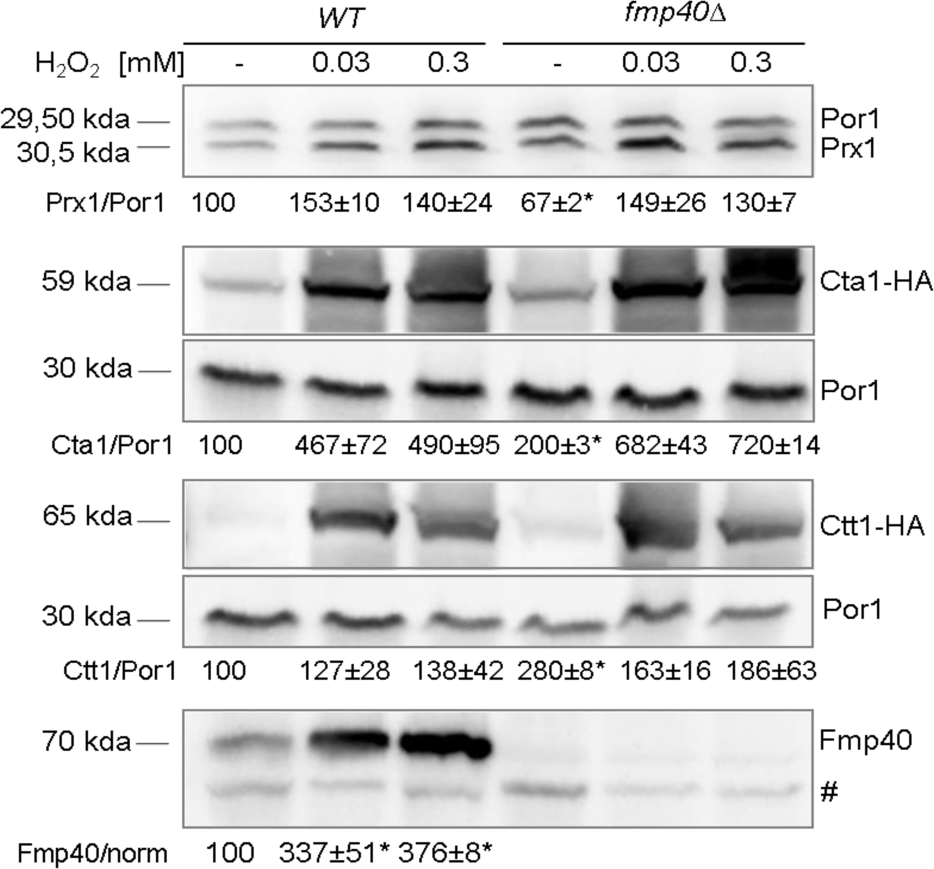
Fmp40, Prx,1 Cta1, and Ctt1 protein levels in *fmp40Δ* cells in response to growing concentration of hydrogen peroxide. The level of Prx1, Cta1-HA, and Ctt1-HA (3’ genomic fusion with the 3xHA tag encoding sequence in the wild type and *fmp40Δ* strains, respectively) upon exposition to indicated concentrations of hydrogen peroxide. Strains were grown as described in the legend of Figure 3. Proteins were precipitated using 10% TCA and analyzed by SDS-PAGE (on 10% gel) and Western blotting. The protein level was determined by densitometry and normalized to the respective Por1 signal. # - unspecific band recognized by anti-Fmp40 antibody. The statistical significance of the changes in *fmp40Δ* cells in comparison to the control cells was quantified with the unpaired student t-test and is indicated by the *, where the p-value was less than 0.05. The representative blots are shown.

### 3.5. The genetic interactions of FMP40 with PRX1, TRX3, and GRX2, genes encoding mitochondrial redoxins

Since Fmp40 is a mitochondrial protein, we limited further experiments to the investigation of Fmp40 protein function in the regulation of mitochondrial redoxins Prx1, Trx3, and Grx2. We constructed the strains lacking genes encoding these redoxins (single mutants) and strains lacking both the gene encoding selected redoxin and the *FMP40* gene (double mutants). We studied the influence of respective deletions on the growth phenotypes of tested mutants on a solid, rich glucose medium in the presence of the growing concentrations of hydrogen peroxide. The *prx1Δ* and *trx3Δ* strains conferred slight resistance to hydrogen peroxide concentrations, while the lack of *GRX2* did not affect the strain growth in the applied conditions (Fig. 5A). Lack of Fmp40 increased the hydrogen peroxide sensitivity of *prx1Δ*, *grx2Δ* and *trx3Δ* mutants; the *grx2Δ fmp40Δ* cells were the most sensitive. The results showed that the resistance of *prx1Δ*, *trx3Δ,* and *grx2Δ* strains to the high hydrogen peroxide concentration depends on Fmp40 presence.

**Fig. 5.**
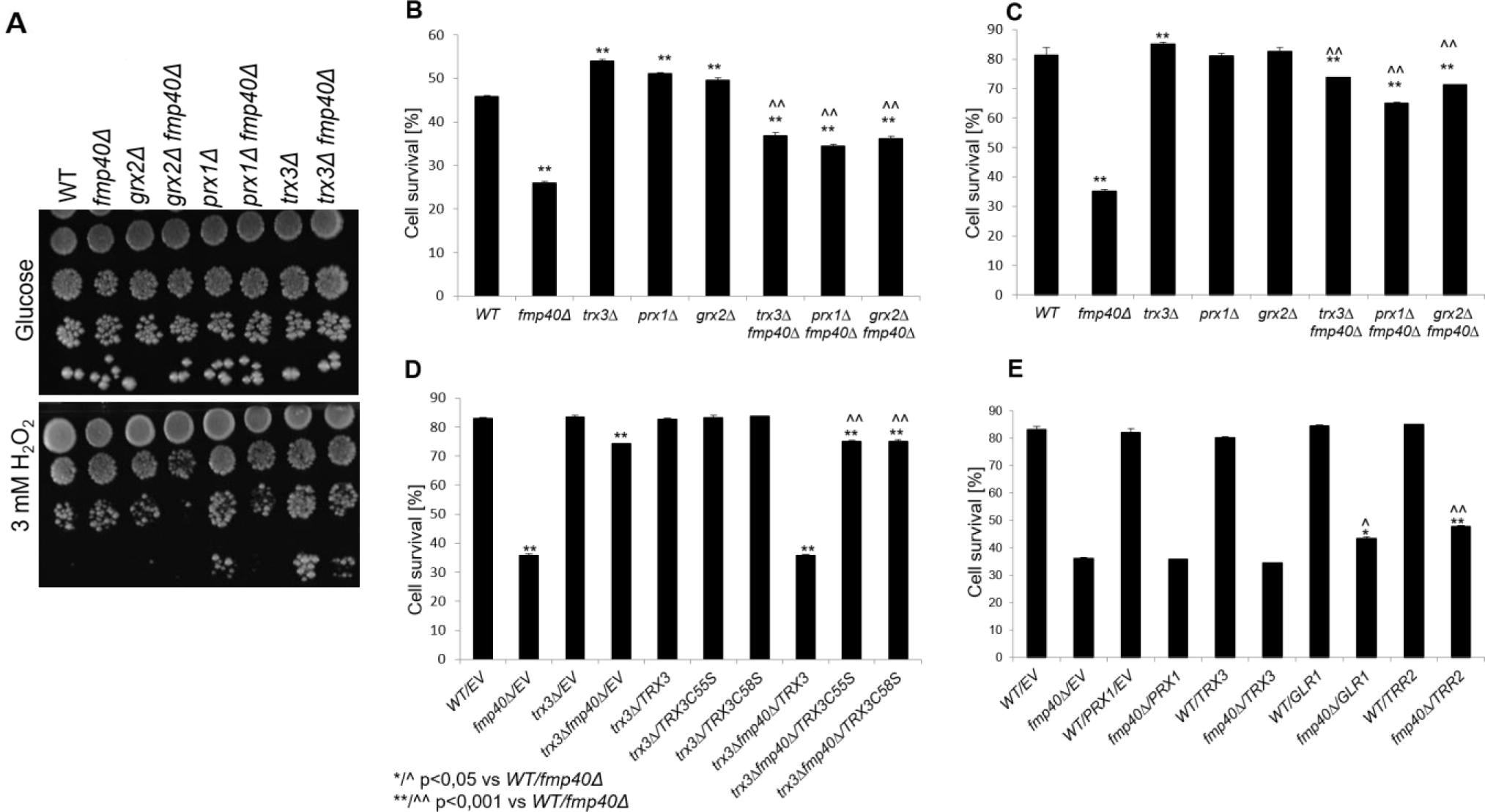
The decreased survival of *fmp40Δ* cells in the presence of hydrogen peroxide depends on mitochondrial redoxins, especially the oxidation of Trx3. (A) Cells pre-grown o/n in rich glucose medium were serially diluted and spotted on YPGluA solid medium without or with a given concentration of hydrogen peroxide, incubated at 28 °C for three days. Cells grown o/n in rich glycerol **(B, C)** or minimal glucose medium **(D, E)** were exposed to 0.03 mM **(B, C)** or 0.1 mM H_2_O_2_ **(D, E)** for 200 minutes, and then the serial dilutions were plated on rich glucose medium and incubated two days at 28 °C. The colonies were counted and cell viability was then determined considering the number of colonies grown from non-treated culture as 100 %. The unpaired student t-test was used to quantify the statistical significance. Indications mean: */^ p<0.05, **/^^ p<0.01 vs WT / *fmp40Δ* cells, EV – empty vector.

We further explored the lower survival of *fmp40Δ* cells observed in CFU experiment after treatment with lower hydrogen peroxide concentrations since such conditions permitted receiving quantitative results for the sensitivity phenotype. Data showed that *fmp40Δ* sensitivity observed after treatment with lower concentrations of hydrogen peroxide could be reversed by removal of the mitochondrial redoxins Trx3, Prx1, or Grx2 from the cells (Fig. 5B, C). Since the oxidized form of Trx3 was shown to promote programmed cell death in yeast ^38^, we asked whether the Trx3 variant that cannot be oxidized, due to the substitution of Cys_55_ to Ser or Cys_58_ to Ser could reverse the *fmp40Δ* cells sensitivity phenotype as native Trx3 protein. The complementation assay revealed that while introduction of vector expressing *TRX3* gene to the *trx3Δ fmp40Δ* cells sensitizes them to H_2_O_2_ to the level typical for *fmp40Δ* cells, the introduction of vector expressing *trx3-C_55_S* or *trx3-C_58_S* does not (Fig. 5D). Thus, oxidation of respective cysteine residues in Trx3 is crucial for its Fmp40-dependent function during oxidative stress. The overexpression assays using the other redox proteins encoding genes revealed that neither overexpression of *PRX1* nor *TRX3* complemented the *fmp40Δ* cells sensitivity phenotype; however, overexpression of *TRR2* or *GLR1* slightly but significantly improved *fmp40Δ* cells survival in low oxidative stress conditions (Fig. 5E). The decreased survival of *fmp40Δ* cells upon treatment with lower hydrogen peroxide concentration was also reversed by the lack of Yca1, a protease involved in programmed cell death (Fig. S3C). We conclude that *fmp40Δ* cells, after exposition to hydrogen peroxide, die in Yca1 and oxidized-Trx3-form-dependent fashion. However, this effect also depends on the stressor concentration, as cells react differently to low and high H_2_O_2_ concentrations. We also assume Fmp40 to participate in ROS control mechanisms in multiple ways because the overproduction of each from tested general reductases, Glr1 and Trr2, improved the viability of *fmp40Δ* cells in the stress conditions.

### 3.6. Fmp40 is necessary to maintain proper NADPH, GSH, and GSSG pool

Since, it was found that Fmp40 interacts with mitochondrial redoxins Prx1, Trx3, and Grx2 whose activity depends on reducing force provided by glutathione and/or NADPH, and glutathionylation of proteins in mitochondria upon diamide exposition was lower in cells lacking Fmp40 ^10^, we tested whether lack of Fmp40 might affect the pools of GSH and NADP(H). Deletion of *FMP40* significantly increased or decreased the glutathione and NADPH pools in yeast cells depending on experimental conditions (presence or absence of H_2_O_2_). In control conditions in *fmp40Δ* cells, GSH level and GSH/GSSG ratio were increased (Fig. 6A), while NADPH level and NADPH/NADP^+^ ratio decreased (Fig. 6B). After treatment with hydrogen peroxide in *fmp40*Δ cells, a significant drop in GSH level and GSH/GSSG ratio was observed (Fig. 6A). Worth to note is that the level of oxidized glutathione (GSSG) in *fmp40*Δ was not changed between conditions with or without hydrogen peroxide. The lowest or not changed GSSG levels were also observed in the *prx1Δ* and *trx3Δ fmp40Δ* mutants (in the stress conditions only). In general (particularly noticeable in control conditions), deletion of *GRX2, PRX1*, or *TRX3* in *fmp40*Δ cells restored the GSH level and GSH/GSSG ratio to those noted in the WT strain. In turn, simultaneous deletion of *PRX1* and *FMP40* in cells exposed to hydrogen peroxide resulted in a significantly decreased level of GSH and GSH/GSSG ratio with relatively high levels of GSSG (Fig. 6A). Subsequently we explored the changes in cellular NADP(H) pools upon treatment with hydrogen peroxide. In general, we have seen that cells respond to hydrogen peroxide by significantly increasing NADPH levels and NADPH/NADP^+^ ratio, which suggests the increased demands for NADPH-based reducing systems. The highest NADPH levels, significantly higher than typical for the wild-type strain, were detected in *prx1Δ* and *trx3Δ* mutants. High NADPH levels characteristic for these mutants were *FMP40*-independent, as no reduction of their levels was found in respective double mutants lacking the *FMP40* gene. However, in the case of *trx3Δ* and *trx3Δ fmp40Δ* mutants, there was also increased NADP^+^ content, which resulted in a decreased NADPH/NADP^+^ ratio after hydrogen peroxide treatment in comparison to the control strain. Thus, a significantly increased NADPH/NADP^+^ ratio was noted only in the case of *prx1Δ* and *prx1Δ fmp40Δ* mutants (Fig. 6B). The obtained results correspond to mutants growth phenotypes and complementation of decreased survival of *fmp40Δ* cells by overexpression of Glr1 or Trr2 (Fig. 5). We conclude that preventing the increase in oxidized glutathione level (through various routes) is a possible way to increase the survival of cells exposed to hydrogen peroxide. Moreover, glutathione and NADP(H) systems work together but also counterbalance each other to keep cellular redox balance, and a high level of NADPH can be used by cellular reductases to reduce GSSG.

**Fig. 6.**
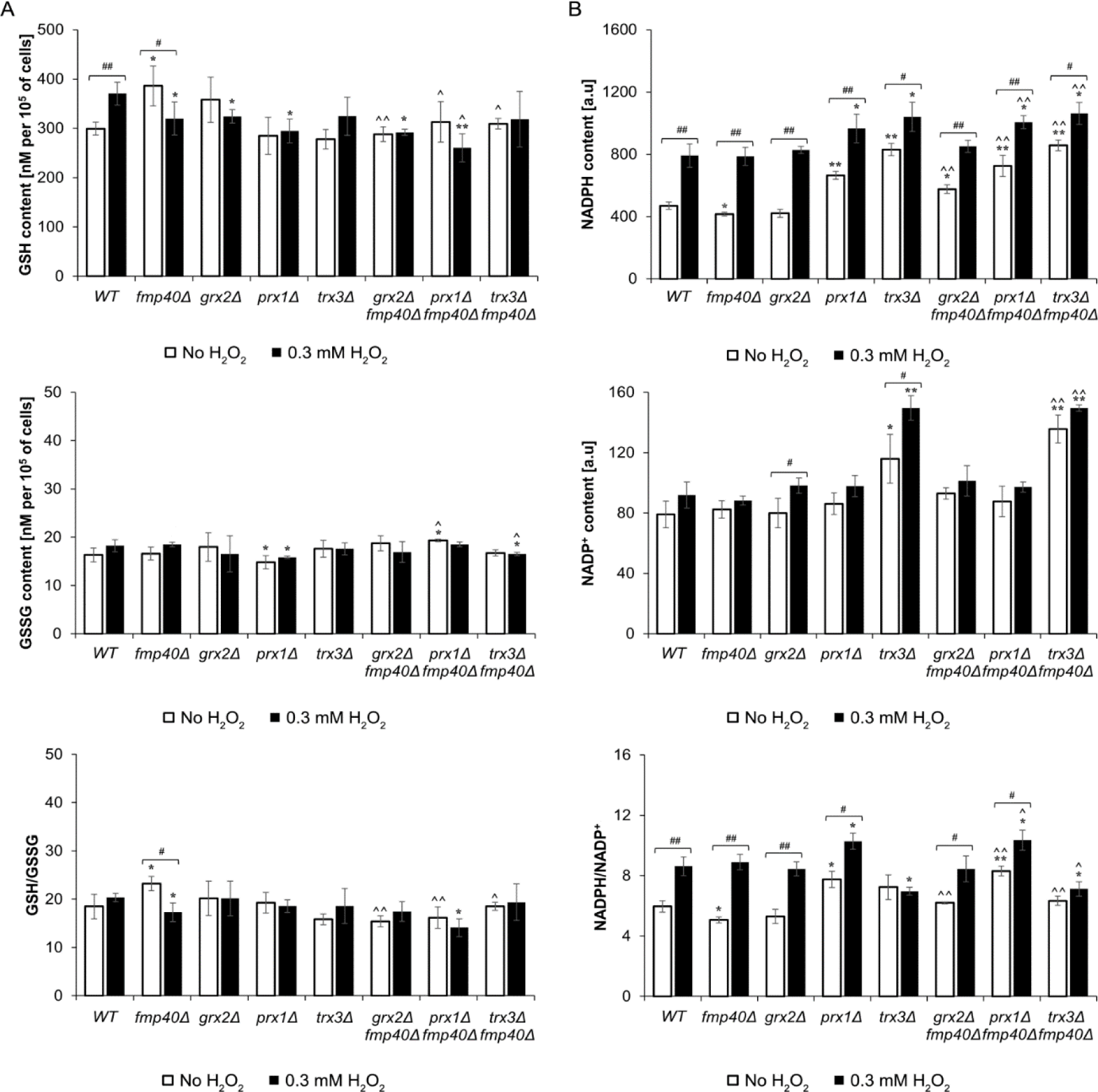
The survival of cells exposed to hydrogen peroxide depends on NADP(H) balance and reduction of oxidized glutathione. **(A)** The glutathione content and GSH/GSSG ratio in the presence or absence of H_2_O_2_. **(B)** The NADP(H) content and NADPH/NADP^+^ ratio in the presence or absence of H_2_O_2_. Strains were grown in YPGlyA medium and subsequently treated with 0.3 mM H_2_O_2_ for 200 min. Glutathione and NADP(H) contents were determined using the luciferin/luciferase-based luminescence method (see Material and Methods section for details). The results are presented as mean ± SD from at least three independent experiments. The statistical analysis was performed using the STATISTICA 13.3 software. The statistical significance of the differences in comparison to the WT/*fmp40*Δ strain was evaluated using one-way ANOVA with the Tukey *post-hoc* test. The statistical significance of the differences between values obtained with or without H_2_O_2_ was evaluated using the t-test for independent samples. The values were considered significant at p-value < 0.05. Used designations: */^ p < 0.05, **/^^ p < 0.001 vs WT/*fmp40*Δ, # p < 0.05, ## p < 0.001 comparing cells grown with or without H_2_O_2._

### 3.7. The Prx1 is not efficiently reduced when Fmp40 is absent

Given that the phenotype of *fmp40Δ* cells depends on the oxidation state of Trx3, we tested whether Fmp40 might regulate the Trx3 redox state. The oxidation state of proteins can be preserved by rapid treatment of cells with trichloracetic acid (TCA), which protonates free thiol groups. Afterward, the protein extracts were reacted with the thiol-specific probe 4-acetamido-4’maleimidyldystilbene-2,2’-disulfonic acid (AMS). AMS alkylates cysteine residues in a free-SH, but not in an oxidized state, increasing their relative molecular mass by 0.5 kDa, which SDS-PAGE / Western blot analysis allows to detect if only analyzed protein is small enough ^13^. The experiment was performed with the myc-tagged Trx3 protein (expressed from the genomic *TRX3* locus) in the *fmp40Δ*, *prx1Δ*, and *grx2Δ* single deletion mutants as well as the double *fmp40Δ prx1Δ* and *fmp40Δ grx2Δ* mutants, without and with exposition to H_2_O_2_, as in previously described experiments. Samples were either (i) treated with the reducing agent Tris(2-carboxyethyl)phosphine (TCEP), (ii) treated with AMS, or (iii) treated with both TCEP and AMS (reduced). TCEP reduces disulfide bonds and sulfenic acids but not sulfinic or sulfonic acids protein derivatives ^63^.

In the non-stressed cells, the migration of Trx3 in all samples isolated from the wild-type, *fmp40Δ*, *prx1Δ*, and *fmp40Δ prx1Δ* strains grown under respiratory conditions was decreased following treatment with AMS, indicating that the vast majority of Trx3 is present in the reduced form in these conditions (Fig. 7A and S4A). Loss of *GRX2* shifted the Trx3 redox balance to an oxidized state and a small amount of oxidized form of Trx3 accumulated, similarly in the single *grx2Δ* and the double *fmp40Δ grx2Δ* cells untreated and treated with hydrogen peroxide. Moreover, after exposition to 0.3 mM H_2_O_2_, this form was present in the samples treated with TCEP, indicating that it is irreversibly oxidized. Thus Grx2 protein is indeed involved in the reduction of Trx3 *in vivo*, as suggested previously based on the *in vitro* experiments ^35^. After exposition to 0.03 mM H_2_O_2_ a small amount of oxidized Trx3 form is present in *fmp40Δ* cells but not in the control cells, in which this form appears only after exposition to 0.3 mM H_2_O_2_. Interestingly, in these conditions, the oxidized form of Trx3 is less abundant in *fmp40Δ* cells, which is consistent with the different survival of *fmp40Δ* cells upon treatment with H_2_O_2_.

**Fig. 7.**
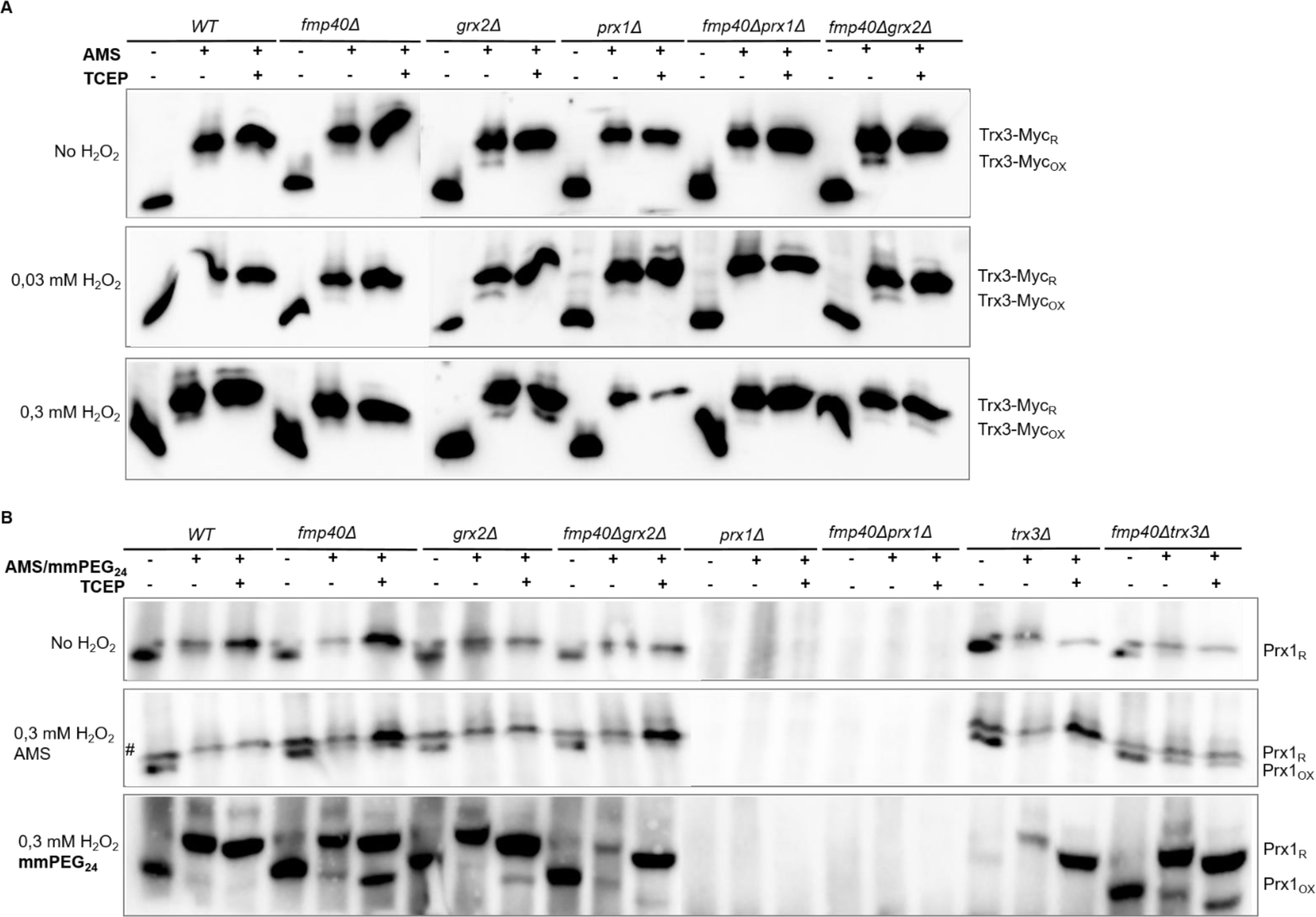
Redox state of the Trx3 and Prx1 proteins, according to the redox shift assay. **(A)** The *TRX3-Myc* fusion gene was introduced to the yeast genome in the indicated strains, which allowed its expression from the native *TRX3* promoter. Strains were grown to the exponential phase in the YPGlyA medium and treated with the indicated concentrations of H_2_O_2_ for 200 min. Proteins were extracted, incubated with AMS or TCEP as displayed, separated in 18% SDS-PAGE gels, transferred to the membrane, and detected with anti-Myc antibody. **(B)** The indicated strains were grown as above, proteins were extracted, and incubated with TCEP or AMS / mmPEG_24_ as indicated. Proteins were analyzed in 12% (AMS) or 14% (mmPEG_24_) SDS-PAGE respectively. After transfer to the membrane, the Prx1 protein was detected using an anti-Prx1 antibody. The nature of the Prx1-antibody decorated band migrating slower in AMS-untreated samples, marked with #, is unknown. The representative blots are shown.

We then asked about the redox state of Prx1 mitochondrial peroxiredoxin. To observe the SDS– PAGE-based redox shift of Prx1 we used both AMS and the maleimide compound mmPEG_24_, which upon modification of a cysteine thiol results in an increased mass of 2.5 kDa ^50^. The experiment was performed in *fmp40Δ*, *prx1Δ, trx3Δ*, and *grx2Δ* single deletion mutants as well as the double *fmp40Δ prx1Δ, fmp40Δ trx3Δ*, and *fmp40Δ grx2Δ* mutants, without and with exposition to 0.3 mM H_2_O_2_. Upon exposition to the high concentration of hydrogen peroxide, the appearance of an irreversibly oxidized form of Prx1 is visible in all the strains lacking *FMP40* (Fig. 7B and S4A). Thus, the Fmp40 is critical for the efficient reduction of Prx1 during high oxidative stress conditions. To some extent, the lack of Grx2 also impacts the reduction of Prx1, because the oxidized form of Prx1 is visible in AMS/TCEP-treated samples from cells lacking Grx2, although it is not visible in samples treated with only AMS or only mmPEG_24_. This is due to the reduced protein stability in samples treated with these compounds without the presence of TCEP, previously observed by other groups ^26^. A similar experiment was performed looking for the Grx2 redox state (Fig. S4B). The study of Grx2 redox state is difficult due to several forms of this protein present in cells: cytosolic and mitochondrial forms, including the precursor form ^24,64^. Despite these caveats, we performed such an experiment, which showed no particular changes in the Grx2 protein migration pattern in the mutant strains when compared to the wild-type strain.

### 3.8. Fmp40 AMPylates redoxins Prx1, Trx3 and Grx2 in vitro and interacts with Trx3-Myc in vivo

To uncover how Fmp40 regulates the redox state of Trx3 and Prx1, we first asked whether Fmp40 transfers AMP to Prx1, Trx3, and Grx2 in the *in vitro* assay. To answer this question, we purified Prx1, Trx3, and Grx2 from bacteria; then, the purified proteins were incubated with or without Fmp40 and with [α^32^P]ATP. We included the cytosolic Tsa2 peroxiredoxin in the experiment as a negative control, expecting that it would not be AMPylated by the mitochondrial AMPylase ^65^. The auto-AMPylation of Fmp40 was used as a positive control in this experiment ^10^. We found the ^32^P-incorporation to the redoxins Prx1, Trx3 and Grx2 but not Tsa2 protein, and in the presence of Fmp40 only (Fig. 8A).

To indicate the modified amino acid residues, the reactions with Prx1, Trx3, Grx2, and Fmp40 were repeated with unlabeled ATP and the reaction products were analyzed by mass spectrometry. Among peptides resulting from trypsin digestion, we identified modification by AMP on Ser_116_ of Prx1, Thr_95_ of Trx3, and Thr_32_, Thr_108_ and/or Ser_123_ of Grx2 (Fig. S5A). We introduced mutations into the coding sequence of *PRX1* and *TRX3* genes, cloned into the pET15b vector, to introduce the alanine substitutions of the amino acid residues indicated as the subject of AMPylation in the mass spectroscopy approach. The substituted variants of the proteins were produced and ampylation assays *in vitro* were repeated. We still could observe the ^32^P incorporation to the Prx1-S_116_>A and Trx3-T_95_>A protein variants after AMPylation reaction *in vitro* (Fig. S5B), suggesting that the modification can be held in the other residues. It should be mentioned here that Prx1 and Trx3 contain, respectively: 17 and 5 serine, 5 and 4 tyrosine and 18 and 9 threonine residues. Aiming to find the modified residues in the Trx3 protein in the *in vivo* experiment, we used the wild-type and *fmp40Δ* strains currying the *TRX3-Myc* gene in their genomes and the α-Myc-coated magnetic beads to purify the Trx3-Myc fusion protein from freshly isolated mitochondria. Then, we performed the comparative mass spectrometry analysis. With this approach, the threonine in position 66 of Trx3 was found AMPylated (Fig. S6).

In the similarly performed experiments, but analyzed by Western blotting, we looked for proteins that co-purified with Trx3-Myc and found that about 15 % of the total Fmp40 and about 3,6 % of the total Prx1 pools in the beads eluates (Fig. 8B). The physical interaction between Fmp40 and Trx3, which is a mitochondrial matrix protein ^42^, suggested, that Fmp40, in addition to its confirmed localization in the mitochondrial intermembrane space ^66^, may have a matrix echoform. To check this possibility, we used the BiG Mito-Split-GFP yeast strain, expressing the non-fluorescent GFP-β1-10 fragment from the mitochondrial genome, previously created in our team for the identification of matrix proteins ^67^. The gene encoding Fmp40 fused to a GFP-β11 fragment, cloned into a centromeric plasmid, when introduced to this strain, gave a clear fluorescence signal, providing proof of the Fmp40 presence in the mitochondrial matrix (Fig. S7). Therefore, Fmp40, similarly to Prx1 ^33^, has a localization in both the matrix and IMS of mitochondria.

As was mentioned above, the mass spectrometry approach allowed the identification of amino acid residue in Trx3 that undergoes AMPylation, namely Thr_66_. To investigate the biological significance of this modification of the Trx3 protein, we introduced mutations into the codon 66 of the *TRX3* gene in the genomic locus, resulting in respective amino acid substitutions, one threonine (T) to alanine (A), second threonine to glutamic acid (E). Both variants were introduced to two genetic backgrounds of *S. cerevisiae*, MR6 and BY4741. We examined the effect of the Trx3-T_66_>A and Trx3-T_66_>E variants on growth on solid and liquid media with glucose and glycerol, and in the presence of hydrogen peroxide (on solid medium only) at two temperatures 28 °C and 36 °C (the conditions of moderate heat stress for yeast, where growth phenotypes are often more pronounced). The growth of analyzed strains in the tested conditions did not differ significantly (Fig. S8). However, in the MR6 background, at an elevated temperature, on both carbon sources, cells expressing the Trx3-T66>E variant later than wild-type cells stop growing reaching the stationary phase.

To examine the level of Trx3, Trx3-T_66_>A, and Trx3-T_66_>E proteins, we introduced a sequence encoding the triple HA tag at the 3’ end of the *TRX3* gene in these strains, in both the wild type and *fmp40Δ*, obtaining a recuiredset of strains in the MR6 strain background. We examined the level of Trx3 variants in cells grown at 28 °C on complete medium with glycerol, without and after exposure to hydrogen peroxide. The level of Trx3 protein in cells not exposed to hydrogen peroxide is low and increases twofold and fourfold, when cells are exposed to 0.03 and 0.3 mM hydrogen peroxide, respectively (Fig. 8C). The levels of Trx3-T_66_>A and Trx3-T_66_>E increase as well but to a lower extent, still with statistical significance, in cells exposed to 0.03 mM hydrogen peroxide. Reduction in wild-type Trx3 accumulation upon 0.3 mM hydrogen peroxide treatment is also seen in the *fmp40Δ* mutant. The Trx3 protein has a mitochondria-targeting sequence within its first 21 amino acids ^42^. Upon hydrogen peroxide treatment, the precursor form of the Trx3 protein accumulates, and its amount increases in cells exposed to 0.3 mM hydrogen peroxide to 28%. The precursor form of the Trx3-T_66_>A and Trx3-T_66_>E variants is more abundant – 41 and 51%, respectively. Interestingly, the amount of the unprocessed form of Trx3 protein increases in the *fmp40Δ* mutant to the same extent in stressed cells. To conclude, the T_66_ in Trx3 redoxin is important for the regulation of its functional level upon oxidative stress conditions.

## 4. Discussion

### Fmp40 function is to control the cell’s response to oxidative stress and survival

The survival of yeast cells depends on how they cope with stress. Many stress conditions leading to ROS production, including H_2_O_2_ exposure and depletion of GSH, have been shown to promote yeast PCD ^68,69^. Cellular respiration and oxidative phosphorylation in mitochondria are accompanied by the production of ROS, which play a role in signal transduction ^70–73^. Several mechanisms influence the maintenance of an appropriate level of ROS, and the expression of many redox proteins, including mitochondrial redoxins, is induced under conditions of cellular respiration, e.g., growth of yeast cells on non-fermentable carbon sources like glycerol ^74,75^. Fmp40 is one of redox proteins and its absence leads to dysregulation of other redox genes expression. In *fmp40Δ* background expression of general *MSN2*, *SKN7*, and *YAP1* transcription factors involved in oxidative stress response showed a significant decrease. The expression of the *SOD1* gene, encoding a bifunctional protein (the superoxide dismutase and transcription factor regulating the expression of oxidative resistance and repair genes ^76^) showed different behavior. Its expression increased significantly with the growing concentration of hydrogen peroxide; however, the initial level of *SOD1* mRNA in the *fmp40Δ* strain was considerably lower than in the control strain, so despite a faster increase in expression upon oxidative stress, the *fmp40Δ* to WT *SOD1* expression ratio became doubled only in high oxidative stress conditions. Similar dependence on *fmp40Δ* in the expression pattern showed the group of genes encoding proteins protecting cells against oxidative stress, e.g., *TSA1*, *TRR2*, *GRX8*, but also *TRX3*, *GRX3*, *GTT2*, *GRX2* that have other dynamics of expression in response to the rising concentration of hydrogen peroxide in each of tested strains: decreased at non-stressed cells or treated by 0.03 mM but increased at 0.3 mM hydrogen peroxide treatment. These results highlight the complexity of regulation of the cellular response to oxidative stress in which Fmp40 is an important regulator. It may be different under different growth conditions: depending on the carbon source, temperature, solid or liquid culture, growth phase, stressor dose, or treatment time ^77,78^. The molecular mechanisms of regulation may vary, e.g., a posttranslational modification. In the case of the oxidative stress response, the well-known examples are the phosphorylation of Sod1 or redox state control of Yap1 that influence their DNA binding activity, half-life time, and localization in/out of the nucleus, thus controlling their function as transcription factors ^76,79–81^. Fmp40 is one of the proteins that regulate the cell’s response to oxidative stress through a post-translational modification, such as AMPylation, while also being regulated by the oxidation/reduction state ^10^. Changes in gene expression in *fmp40Δ* cells depending on the hydrogen peroxide concentration correlate with the opposite growth phenotypes we found. The *fmp40Δ* cells have reduced survival upon exposition to low concentrations of hydrogen peroxide in the W303-1B or menadione in BY4741 genetic backgrounds, phenotype dependent on the activity of Yca1 caspase, and therefore on the process of programmed cell death ^82,83^. However, in the BY4741 strain background, in higher hydrogen peroxide concentrations, the lack of Fmp40 resulted in less frequent apoptosis and, thus, increased survival. Differences in sensitivity to various compounds in media that generate cellular stress between these genetic backgrounds of yeast cells are well documented in the literature, where the W303-1B strain is in general more sensitive than BY4741, therefore resistance phenotypes are often not visible in W303-1B background ^84,85^. The differences might be explained partially by the presence of different gene variants or mutant alleles in these strains’ genomes that compromised encoded products’ function and, subsequently, cells’ growth abilities and effectiveness of stress response ^86^. The differences are apparent even on the proteomic level while comparing proteomes of W303- and BY474X-derivatives being at different growth stages (e.g., exponential versus stationary phase) ^87^.

Under conditions of respiratory growth, the functionality of Fmp40, like Grx2, influences GSH consumption. This is supported by a high level of GSH with normal expression of *GSH1* and *GSH2* genes required for GSH synthesis ^88,89^ and reduced expression of enzymes that use GSH: mainly *GRX2, GTT1*, *GTT2, GRX3*, *GRX5*, *GRX8* in *fmp40Δ* cells, determined in conditions without hydrogen peroxide. Reduced glutathione consumption in *fmp40Δ* cells appears to be connected with a previously noted decrease in global S-glutathionylation ^10^ but also with the activity of Trx3, Grx2, and Prx1 function, as lack of these proteins restores the GSH level in *fmp40Δ* cells to that of the control cells. These results suggest that the lack of Fmp40 alters the mitochondrial GSH-dependent reduction processes of Prx1, Trx3, and/or Grx2. This is consistent with data from the literature indicating that Prx1 reduction is a major source of GSSG in mitochondria, the level of which triggers a transcriptional response to upregulate the cytosolic catalase, Ctt1, demonstrating that cells can recognize and respond to disturbed matrix redox homeostasis ^15^. At the same time, the activity of Fmp40 affects NADPH to a much lesser extent than GSH. However, the GSH/GSSG and NADPH/NADP^+^ ratios are changed inversely in non-stressed *fmp40Δ* cells (increased and decreased, respectively), which may signal a change in the expression of redox enzymes genes - generally a decrease in their expression. This results in decreased survival after exposure to low concentrations of hydrogen peroxide in the W303-1B background or to menadione in the BY4741 background ^10^. In accordance with that, the percentage of cells in which the Nuc1 nuclease re-localizes to the nucleus (in non-stressed but respiratory conditions) in the *fmp40Δ* BY4741 strain is increased.

After exposure to high concentrations of hydrogen peroxide the demand for GSH-dependent processes increases, the GSH pool decreases in *fmp40Δ* cells, and therefore the GSH/GSSG ratio, without changes in the NADPH/NADP^+^ ratio, even though one of the compensating mechanisms is the reduction of protein glutathionylation in mitochondria ^10^. The expression of many redox enzymes increases, including those that encode redox enzymes consuming GSH, and cells have better survival. The NADPH, in turn, is consumed in reduction processes dependent on Prx1 and Trx3 - better hydrogen peroxide tolerance in cells without Prx1 or Trx3 correlates with a higher level of NADPH. Fmp40 has less impact on the NADPH/NADP^+^ pools and ratio, but a rescue of the sensitivity to hydrogen peroxide in *pos5Δ* by *fmp40Δ* indicates that Fmp40 impacts NADPH consumption as well ^90^. The uncharacterized Ycp4 mitochondrial flavoredoxin may be involved in these processes as the sensitivity of *ycp4Δ* cells is rescued by *fmp40Δ* to a similar extent as that of *pos5Δ* cells.

The changes in the strains’ sensitivity to hydrogen peroxide revealed the genetic interactions that imply an essential role of Fmp40 in ROS and PCD signaling. The profound synthetic growth defects showed the *grx5Δ* and *sod2Δ* when combined with *fmp40Δ*. The synergistic epistatic relation between *fmp40Δ* and *sod2Δ* pointed out the crucial role of Fmp40 in the hydrogen peroxide neutralization in the mitochondrial matrix, independently from Sod2 ^91^. Following that, is our finding that Fmp40, besides its presence in the IMS, localizes in the mitochondrial matrix. Lack of *GRX5* leads to perturbations in the Fe-S cluster biogenesis^92^. Since it was previously shown that in the *prx1Δ* cells, transcription of Aft1-dependent genes, ensuring iron utilization and homeostasis (including Fe-S clusters formation) was increased, while biogenesis of Fe-S clusters was efficient, we believe that Fmp40 is not involved in Fe-S clusters biogenesis process ^93^. The synthetic growth defect of *fmp40Δ* with *grx5Δ* suggests dysfunction of the mitochondrial signaling to the cytosol, accompanied by labile iron release accelerating cell death caused by lack of Grx5. We could assume it because removing the *FMP40* gene from the *prx1Δ* genome leads to the partial rescue phenotype, e.g., lower sensitivity to hydrogen peroxide of double mutant than in the single *prx1Δ* strain.

Complementation of the *fmp40Δ* cells decreased survival after exposition to low hydrogen peroxide concentration by removal of the *YCA1* gene encoding the homolog of mammalian metacaspase or by replacement of native *TRX3* gene by its variant encoding the non-oxidizable form of Trx3, and variations in the *AIF1* gene expression, are solid arguments stating for the participation of Fmp40 in PCD signaling. The Yca1 cysteine protease is required to induce PCD mediated by oxidized Trx3 ^38^. The GSH/GSSG balance is changed during apoptosis ^8^. The yeast oxidoreductase Aif1, a flavoprotein that localizes in mitochondria (in mammalian cells in IMS ^94^), is the homolog of human apoptosis-inducing factor AIF ^95,96^. In response to pro-apoptotic stimuli Aif1, alike Nuc1, translocates to the nucleus contributing to DNA fragmentation. However, due to its similarity to mammalian AIF, for which it was shown that it has NADH oxidase and ROS scavenging activity in mitochondria, it was proposed that Aif1 plays a physiological role in redox homeostasis as well ^97^. Moreover, mammalian AIF is believed to play a dual role, both pro- and anti-apoptotic, in response to low and high oxidative stress, respectively, similar to what we observe in *fmp40Δ* cells (see ^69^ and references within). We postulate that the opposite phenotypes of *fmp40Δ* cells grown under non-fermentative conditions and exposed to different levels of oxidative stress are due to the loss of control of Prx1 and Trx3 reduction cycles and consequently ROS signaling.

### The model of Fmp40 and Grx2 function in the Prx1 reduction cycle

The literature data concluded that: (i) GSH is part of the reductive mechanism of Prx1 but is not oxidized in the process, forming a disulfide bond with the Prx1 peroxidatic cysteine, which is reduced by Trx3 ^98^; (ii) GSH forms a transient mixed disulfide with the Prx1 peroxidatic cysteine that is subsequently reduced by Trr2, leading to the formation of a transient intermolecular disulfide between Prx1 and Trr2, that is reduced by GSH leading to GSSG formation ^36^; (iii) the mitochondrial thioredoxin system, Trx3, and Trr2, can efficiently reduce Prx1 *in vitro* ^31^; (iv) the Prx1-GSH is reduced by Trx3 with the transient intermolecular disulfide between Prx1 and Trx3 ^38^; (v) Prx1-GSH transient intermediate is reduced by Grx2 *in vitro*, ultimately leading to GSSG formation ^35^. In this work, we showed that Grx2 is involved in the reduction of Trx3 and Prx1 *in vivo* while Fmp40 impacts mainly the reduction of Prx1 in high-stress conditions and reduction of Trx3 in different ways depending on the stress level. In the reduction steps of Prx1, the Grx2 may act at the level of Trx3 reduction and/or Prx1-GSH intermediate reduction (Fig. 9).

**Fig. 9.**
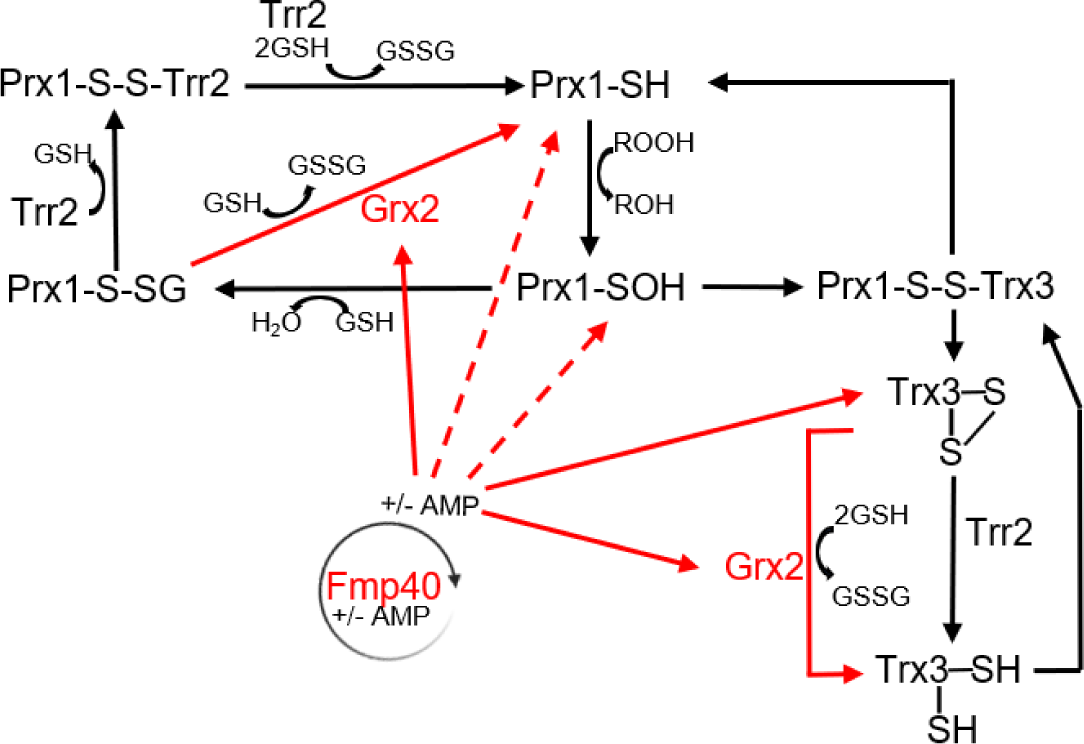
Fmp40 plays an essential role in the Prx1 reduction. By regulating Grx2 activity, Fmp40 impacts the reduction of Trx3 and Prx1, the latter upon high oxidative stress. The molecular mechanisms of regulation by Fmp40 likely rely on the ampylation reaction of Grx2, Trx3, and Prx1. The presence or absence of ampylation of these enzymes may modulate their activity, reduction process, and interactions between them or with GSH. Red arrows indicate the proposed reactions in the Prx1 reduction process based on the results of this study.

We have previously shown that the reduced Fmp40 is active as an AMPylase, thus Fmp40 must be regulated by and/or sense the ROS level ^10^. We propose that under high oxidative stress, Fmp40 undergoes oxidation, which inhibits its AMPylase activity and/or changes its specificity towards substrates. Our results show that the Prx1, Trx3, and Grx2 proteins can be AMPylated by Fmp40, which influence their activity, and/or reduction cycle. Basing on the affinity of Fmp40 to Grx2 in the *in vitro* AMPylation reaction, additive growth defect, and similar changes in the GSH/GSSG and NADPH/NADP^+^ ratios in the single *fmp40Δ* and *grx2Δ* mutants we argue that Grx2, like the *E. coli* GrxA, is the main substrate of Fmp40 ^10^. The second protein on which Fmp40 performs its regulatory function is Trx3, in which substitutions of the Thr_66_ that block or mimic its AMPylation, in conditions of lower oxidative stress, impact the Trx3 protein levels, while in conditions of high stress, inhibit its maturation (like the lack of Fmp40). The direct Fmp40 action on Prx1 may not be excluded (Fig. 9). As AMPylation and phosphorylation take place on the same residues, we postulate that both modifications, involving the same residue, may be responsible for the regulation of the redoxins activity or reduction cycle. It was proposed that both protein kinases and phosphatases would be directly regulated by oxidants, resulting in the autophosphorylation and activation of protein kinases ^99^ and reversible inactivation of protein phosphatases ^100^. No phosphorylated residue has been identified in Trx3 to date, four residues were found phosphorylated in Prx1 while seven were in the Grx2 redoxin. Those are Prx1-S_53_, -S_44_, -S_170_, -S_201_ ^101–103^ and Grx2-S_37_, -T_67_, -S_69_, -S_91_, -S_94_, -S_123_, -T_127_ ^101,104–106^. The biological role of these modifications is not known yet. We hypothesize that different modifications of the same residue by various enzymes may fulfill different roles in redoxins activity regulation. In the case of Trx3, where both T_66_ substitutions to alanine and glutamic acid have the same effect on Trx3 levels and maturation as the absence of Fmp40, it is possible that phosphorylation at this residue may take place in the absence of AMPylase, and also result in the same effect as the absence of AMPylation. To understand the full mechanism of Trx3 regulation, further studies are needed, where the possibility of phosphorylation at this residue should not be excluded. The picture is certainly much more complex, as many PTM are on the cysteine residues and may directly block their redox function. An example of such PTM is palmitoylation and such a regulation of redox activity of mammalian PRDX5 was demonstrated ^107^. In yeast mitochondria, Ycp4 is a palmitoylated flavodoxin-like protein and may undergo a similar regulation ^108^. The suppression by *fmp40Δ* the hydrogen peroxide sensitivity of *ycp4Δ* strain to the level typical for *fmp40Δ* single mutant suggests that Ycp4 works in the same biological process as Fmp40. And indeed, the Ycp4 ortholog of *Escherichia coli*, WrbA, and *Candida albicans* Ycp4, which binds FMN, have NAD(P)H: quinone–oxidoreductase activity. Moreover, Ycp4 is indirectly involved in the regulation of gene expression of metabolic pathways (including sulfate assimilation) and oxidative stress response ^109–111^.

### Mechanisms of PCD inhibition/induction in *fmp40Δ* cells

With exposure of wild-type cells to low hydrogen peroxide concentrations, the Trx3 is activated in the reduction of Prx1, the GSH/GSSH and NADPH/NADP^+^ ratios are kept in balance due to adaptive response by transcription of several antioxidant enzymes mediated by the redox-sensing transcription factors, e.g., Yap1 ^81^. In the absence of Fmp40, due to the ineffective reduction of Trx3, the balance of GSH/GSSG and NADPH/NADP^+^ is changed, and the Trx3-S-S accumulates and stimulates PCD ^38,112^. It is important to note that maintaining the Trx3 in a reduced form is crucial, as it is unaffected even by the loss of Trr2. However, the oxidized mitochondrial Trx3 accumulates in mutants simultaneously lacking both the Trr2 and the glutathione reductase Glr1 ^26^. Thus, the redox state of mitochondrial Trx3 is buffered by the GSH/GSSG balance which depends on the Fmp40. In the wild-type and *fmp40Δ* strains treated with low hydrogen peroxide concentrations, the Aif1 oxidoreductase encoding gene’s expression increases, affecting the expression of many genes encoding proteins necessary for ROS neutralization, or signaling, including apoptosis induction. This changed expression influenced cells’ ability to respond to stress. The direct role of Fmp40 in gene expression may not be excluded, e.g., by AMPylation of a mitochondrial PCD effector (Fig. S9).

At high concentrations of hydrogen peroxide, the Prx1 cellular pool is rapidly hyperoxidized, so it neither degrades hydrogen peroxide nor signals the oxidation of ROS effectors. In *fmp40Δ* cells, hyperoxidation of Prx1 leads to targeting the reduced Trx3 to other substrates, whose redox state influences the transcriptional changes, allowing adaptation to the stress conditions, enabling the repair of oxidative damage, which is necessary for cell survival, as it was suggested previously ^15,113,114^. In parallel, the cellular pool of GSH decreases without changes in the level of GSSG, which indicates GSH consumption for the reduction processes other than Prx1 reduction ^19^. The reduced level of global protein glutathionylation in diamide-treated mitochondria (generates high oxidative stress) in the *fmp40Δ* strain ^10^ indicates that there are mechanisms that inhibit GSH consumption in the process of protein glutathionylation. The redox stress results in higher expression of many genes encoding redox enzymes but a decrease of *AIF1,* which results in effective removal of ROS and higher cell survival.

## Conclusions

Fmp40 is an AMPylase located in the mitochondrial matrix and intermembrane space, whose activity is regulated by its oxidation state. Fmp40 AMPylase is involved in the control of the reduction cycle of mitochondrial redoxins Prx1 and Trx3. Throughout introducing the post-translational modifications to redox proteins, Fmp40 contributes to the ROS signaling during respiration-dependent growth. In addition, Fmp40 is necessary to maintain the balance of cellular redox buffers GSH and NADPH, which also have signaling functions in the cell, as hydrogen peroxide. Fmp40 is necessary for hydrogen peroxide-mediated regulation of the expression of oxidative stress response genes, including those encoding redox enzymes.

Since 22 mutations in the *SELENOO* gene, encoding the human SelO protein, have been identified in patients with congenital genetic diseases (https://www.clinicalgenome.org/data-sharing/clinvar/), understanding the function of SelO proteins is crucial. Further research should clarify how Fmp40 is regulated and indicate the mitochondrial redoxin that modulates the Fmp40 activity. By identifying cellular substrates of Fmp40 AMPylase, it would be possible to answer whether Fmp40 is directly or indirectly involved in the cells’ response to oxidative stress at the transcription level, regulating the expression of genes encoding redox proteins. Finally, considering that AMPylation may compete with phosphorylation of the same amino acid residue, the understanding of controlling the balance between these two modifications in changing environmental conditions might be one of the challenges in future research concerning maintaining redox homeostasis.

## Acknowledgements

Authors thank to Prof. Chris Grant (University of Manchester, UK) for providing strain encoding Trx3-Myc and plasmids encoding the wild type and Trx3-cysteines substitutions variants, Prx1, Glr1 and Trr2. We thank to Prof. Antonio Barcena (University of Jaén, Spain) for anti-Prx1 antibody and pET15b-Prx1, pET15b-Trx3, pET15b-Tsa2 and pET15b-Grx2 expression plasmids, dr Aneta-Kaniak-Golik (Institute of Biochemistry and Biophysics, Polish Academy of Sciences, Poland) for sharing pNUC1-eGFP plasmid, dr Ulrike Topf (Institute of Biochemistry and Biophysics, Polish Academy of Sciences, Poland) for sharing materials and helpful discussions. Special thanks to dr Bruno Senger and prof Hubert Becker (University of Strasburg) for the construction of the pAG414pGPDβ11 expressing Fmp40-β11. We also thank to dr Jakub Piatkowski from the Warsaw University for help in size-exclusion Prx1, Trx3, Grx2, and Tsa2 proteins purification, and dr Renata Zadrąg-Tęcza (University of Rzeszów, Poland) for accepting Suchismita Masanta for a three-week internship to perform GSH/GSSG and NADPH/NADP measurements. We thank A. Malinowska from a Laboratory of Mass Spectrometry, Institute of Biochemistry and Biophysics, Polish Academy of Sciences) for mass spectrometry assistance.

## Statement of Ethics

The permission number for work with genetically modified microorganisms (GMM I) for RK is 01.2-28/201.

## Availability of data and material

Supplementary data and the materials are available upon request.

## CRediT authorship contribution statement

Suchismita Masanta: Investigation, Visualization, Methodology. Aneta Wiesyk: Validation, Methodology, Investigation, Supervision, Resources, Project administration. Chiranjit Panja: Investigation, review & editing. Sylwia Pilch: Investigation. Jaroslaw Ciesla: Investigation. Marta Kasabula: Investigation. Abhipshita De: Investigation. Tuguldur Enkhbaatar: Investigation. Roman Maslanka: Investigation, Supervision, Writing. Adrianna Skoneczna: Writing, review & editing, Visualization, Methodology, Investigation. Roza Kucharczyk: Conceptualization, Writing original draft, review & editing, Supervision, Resources, Project administration, Funding acquisition.

