## SupplementaryData for "Fmp40 ampylase regulates cell survival upon oxidative stress by controlling Prx1 and Trx3 oxidation": MasantaS-SupplementaryFile.pdf

### Abstract

Reactive oxygen species (ROS), play important roles in cellular signaling, nonetheless are toxic at higher concentrations. Cells have many interconnected, overlapped or backup systems to neutralize ROS, but their regulatory mechanisms remain poorly understood. Here, we reveal an essential role for mitochondrial AMPylase Fmp40 from budding yeast in regulating the redox states of mitochondrial 1-Cys peroxiredoxin, Prx1, which is the only protein shown to neutralize $H_2O_2$ with the oxidation of the mitochondrial glutathione and Trx3, thioredoxin, directly involved in the reduction of Prx1. Deletion of *FMP40* impacts a cellular response to $H_2O_2$ treatment that leads to programmed cell death (PCD) induction and an adaptive response involving up or down regulation of genes encoding, among others the catalase Cta1, PCD inducing factor Aif1, and mitochondrial redoxins Trx3 and Grx2. This ultimately perturbs the reduced glutathione and NADPH cellular pools. We further demonstrated that Fmp40 AMPylates Prx1, Trx3, and Grx2 *in vitro* and interacts with Trx3 *in vivo*. AMPylation of the threonine residue 66 in Trx3 is essential for this protein’s proper endogenous level of and its precursor forms’ maturation under oxidative stress conditions. Additionally, we showed the Grx2 involvement in the reduction of Trx3 *in vivo*. Taken together, Fmp40, through control of the reduction of mitochondrial redoxins, regulates the hydrogen peroxide, GSH and NADPH signaling influencing the programmed cell death execution.

### Supplementary Tables

### **Table S1.** Genotypes and sources of yeast strains

| Strain | Nuclear genotype | Source |
|---|---|---|
| W303-1B background | | |
| MR6 | MATa ade2-1 his3-11,15 trp1-1 leu2-3,112 ura3-1 CAN1 arg8::hisG | [1] |
| RKY120 | MATa ade2-1 his3-11,15 trp1-1 leu2-3,112 ura3-1 CAN1 arg8::hisG fmp40::kanMX4 | This study |
| RKY206 | MATα leu2-3,112 trp1-1 ura3-1 ade2-1 his3-11,15 CAN1 arg8::hisG TRX3-MYC::kanMX6 | This study |
| RKY193-2A | MATa leu2-3,112 trp1-1 ura3-1 ade2-1 his3-11,15 CAN1 arg8::hisG fmp40::kanMX4 TRX3-MYC::kanMX6 | This study |
| RKY210-1D | MATα leu2-3,112 trp1-1 ura3-1 ade2-1 his3-11,15 CAN1 arg8::hisG grx2::kanMX4 TRX3-MYC::kanMX6 | This study |
| RKY212-5 | MATa leu2-3,112 trp1-1 ura3-1 ade2-1 his3-11,15 CAN1 arg8::hisG prx1::kanMX4 TRX3-MYC::kanMX6 | This study |
| RKY208-1A | MATa leu2-3,112 trp1-1 ura3-1 ade2-1 his3-11,15 CAN1 arg8::hisG grx2::kanMX4 fmp40::kanMX4 TRX3-MYC::kanMX6 | This study |
| RKY207-3C | MATα leu2-3,112 trp1-1 ura3-1 ade2-1 his3-11,15 CAN1 arg8::hisG prx1::kanMX4 fmp40::kanMX4 TRX3-MYC::kanMX6 | This study |
| SPY3 | MATa ade2-1 his3-11,15 trp1-1 leu2-3,112 ura3-1 CAN1 arg8::hisG grx2::kanMX4 | This study |
| SPY6 | MATa ade2-1 his3-11,15 trp1-1 leu2-3,112 ura3-1 CAN1 arg8::hisG grx2::kanMX4 fmp40::kanMX4 | This study |
| RKY341-5 | MATa ade2-1 his3-11,15 trp1-1 leu2-3,112 ura3-1 CAN1 arg8::hisG prx1::kanMX4 | This study |
| SPY4 | MATα ade2-1 his3-11,15 trp1-1 leu2-3,112 ura3-1 CAN1 arg8::hisG hisG prx1::kanMX4 fmp40::kanMX4 | This study |
| SPY2 | MATa ade2-1 his3-11,15 trp1-1 leu2-3,112 ura3-1 CAN1 arg8::hisG trx3::kanMX4 | This study |
| SPY5 | MATα ade2-1 his3-11,15 trp1-1 leu2-3,112 ura3-1 CAN1 arg8::hisG trx3::kanMX4 fmp40::kanMX4 | This study |
| SPY7 | MATa ade2-1 his3-11,15 trp1-1 leu2-3,112 ura3-1 CAN1 arg8::hisG yca1::kanMX4 | This study |
| SPY8 | MATα ade2-1 his3-11,15 trp1-1 leu2-3,112 ura3-1 CAN1 arg8::hisG yca1::kanMX4 fmp40::kanMX4 | This study |
| RKY358 | Mata ade2-1 his3-11,15 trp1-1 leu2-3,112 ura3-1 CAN1 arg8::HIS3 TRX3-T66>A | This study |
| RKY359 | Mata ade2-1 his3-11,15 trp1-1 leu2-3,112 ura3-1 CAN1 arg8::HIS3 TRX3-T66>E | This study |
| SMY10 | Mata ade2-1 his3-11,15 trp1-1 leu2-3,112 ura3-1 CAN1 arg8::HIS3 TRX3-HA::kanMX6 | This study |
| SMY11 | Mata ade2-1 his3-11,15 trp1-1 leu2-3,112 ura3-1 CAN1 arg8::HIS3 TRX3-T66>A-HA::kanMX6 | This study |
| SMY12 | Mata ade2-1 his3-11,15 trp1-1 leu2-3,112 ura3-1 CAN1 arg8::HIS3 TRX3-T66>E-HA::kanMX6 | This study |
| SMY15 | Mata ade2-1 his3-11,15 trp1-1 leu2-3,112 ura3-1 CAN1 arg8::HIS3 TRX3-HA::kanMX6 fmp40::kanMX4 | This study |
| BiG Mito-Split-GFP | MATa his3-11,15 trp1-1 leu2-3,112 ura3-1 CAN1 arg8::HIS3 [ρ+atp6::GFPβ1-10 5‘UTR$_{COX2}$ ATP6 3‘UTR$_{COX2}$] | [2] |
| BY4741 background | | |
| BY4741 | MAT**a** his3Δ1 leu2Δ0 met15Δ0 ura3Δ0 | Euroscarf |
| BY4742 | MAT**α** his3Δ1 leu2Δ0 lys2Δ0 ura3Δ0 | Euroscarf |
| BY msn2Δ | MATa his3Δ1 leu2Δ0 met15Δ0 ura3Δ0 msn2Δ::kanMX4 | Open Biosystems |
| BY msn4Δ | MATa his3Δ1 leu2Δ0 met15Δ0 ura3Δ0 msn4Δ::kanMX4 | Open Biosystems |
| BY skn7Δ | MATa his3Δ1 leu2Δ0 met15Δ0 ura3Δ0 skn7Δ::kanMX4 | Open Biosystems |
| BY yap1Δ | MATa his3Δ1 leu2Δ0 met15Δ0 ura3Δ0 yap1Δ::kanMX4 | Open Biosystems |
| BY fmp40Δ | MATa his3Δ1 leu2Δ0 met15Δ0 ura3Δ0 fmp40Δ::kanMX4 | Open Biosystems |

| | | |
|---|---|---|
| BY prx1Δ | *MATa his3Δ1 leu2Δ0 met15Δ0 ura3Δ0 prx1Δ::kanMX4* | Open Biosystems |
| BY trx3Δ | *MATa his3Δ1 leu2Δ0 met15Δ0 ura3Δ0 trx3Δ::kanMX4* | Open Biosystems |
| BY grx2Δ | *MATa his3Δ1 leu2Δ0 met15Δ0 ura3Δ0 grx2Δ::kanMX4* | Open Biosystems |
| BY glr1Δ | *MATa his3Δ1 leu2Δ0 met15Δ0 ura3Δ0 glr1Δ::kanMX4* | Open Biosystems |
| BY grx3Δ | *MATa his3Δ1 leu2Δ0 met15Δ0 ura3Δ0 grx3Δ::kanMX4* | Open Biosystems |
| BY grx5Δ | *MATa his3Δ1 leu2Δ0 met15Δ0 ura3Δ0 grx5Δ::kanMX4* | Open Biosystems |
| BY gpx1Δ | *MATa his3Δ1 leu2Δ0 met15Δ0 ura3Δ0 gpx1Δ::kanMX4* | Open Biosystems |
| BY gpx2Δ | *MATa his3Δ1 leu2Δ0 met15Δ0 ura3Δ0 gpx2Δ::kanMX4* | Open Biosystems |
| BY gpx3Δ | *MATa his3Δ1 leu2Δ0 met15Δ0 ura3Δ0 gpx3Δ::kanMX4* | Open Biosystems |
| BY gtt2Δ | *MATa his3Δ1 leu2Δ0 met15Δ0 ura3Δ0 gtt2Δ::kanMX4* | Open Biosystems |
| BY srx1Δ | *MATa his3Δ1 leu2Δ0 met15Δ0 ura3Δ0 srx1Δ::kanMX4* | Open Biosystems |
| BY gsh1Δ | *MATa his3Δ1 leu2Δ0 met15Δ0 ura3Δ0 gsh1Δ::kanMX4* | Open Biosystems |
| BY gsh2Δ | *MATa his3Δ1 leu2Δ0 met15Δ0 ura3Δ0 yap1Δ::kanMX4* | Open Biosystems |
| BY cta1Δ | *MATa his3Δ1 leu2Δ0 met15Δ0 ura3Δ0 cta1Δ::kanMX4* | Open Biosystems |
| BY ctt1Δ | *MATa his3Δ1 leu2Δ0 met15Δ0 ura3Δ0 ctt1Δ::kanMX4* | Open Biosystems |
| BY trr2Δ | *MATa his3Δ1 leu2Δ0 met15Δ0 ura3Δ0 trr2Δ::kanMX4* | Open Biosystems |
| BY trx1Δ | *MATa his3Δ1 leu2Δ0 met15Δ0 ura3Δ0 trx1Δ::kanMX4* | Open Biosystems |
| BY tsa1Δ | *MATa his3Δ1 leu2Δ0 met15Δ0 ura3Δ0 tsa1Δ::kanMX4* | Open Biosystems |
| BY tsa2Δ | *MATa his3Δ1 leu2Δ0 met15Δ0 ura3Δ0 tsa2Δ::kanMX4* | Open Biosystems |
| BY ahp1Δ | *MATa his3Δ1 leu2Δ0 met15Δ0 ura3Δ0 ahp1Δ::kanMX4* | Open Biosystems |
| BY dot5Δ | *MATa his3Δ1 leu2Δ0 met15Δ0 ura3Δ0 dot5Δ::kanMX4* | Open Biosystems |
| BY sod1Δ | *MATa his3Δ1 leu2Δ0 met15Δ0 ura3Δ0 sod1Δ::kanMX4* | Open Biosystems |
| BY sod2Δ | *MATa his3Δ1 leu2Δ0 met15Δ0 ura3Δ0 sod2Δ::kanMX4* | Open Biosystems |
| BY aim6Δ | *MATa his3Δ1 leu2Δ0 met15Δ0 ura3Δ0 aim6Δ::kanMX4* | Open Biosystems |
| BY aim14Δ | *MATa his3Δ1 leu2Δ0 met15Δ0 ura3Δ0 aim14Δ::kanMX4* | Open Biosystems |
| BY aim33Δ | *MATa his3Δ1 leu2Δ0 met15Δ0 ura3Δ0 aim33Δ::kanMX4* | Open Biosystems |
| BY oye2Δ | *MATa his3Δ1 leu2Δ0 met15Δ0 ura3Δ0 oye2Δ::kanMX4* | Open Biosystems |
| BY ycp4Δ | *MATa his3Δ1 leu2Δ0 met15Δ0 ura3Δ0 ycp4Δ::kanMX4* | Open Biosystems |
| BY mdl2Δ | *MATa his3Δ1 leu2Δ0 met15Δ0 ura3Δ0 mdl2Δ::kanMX4* | Open Biosystems |
| BY aif1Δ | *MATa his3Δ1 leu2Δ0 met15Δ0 ura3Δ0 aif1Δ::kanMX4* | Open Biosystems |
| BY mgp12Δ | *MATa his3Δ1 leu2Δ0 met15Δ0 ura3Δ0 mgp12Δ::kanMX4* | Open Biosystems |
| BY oxr1Δ | *MATa his3Δ1 leu2Δ0 met15Δ0 ura3Δ0 oxr1Δ::kanMX4* | Open Biosystems |
| BY pos5Δ | *MATa his3Δ1 leu2Δ0 met15Δ0 ura3Δ0 pos5Δ::kanMX4* | Open Biosystems |

| | | |
|---|---|---|
| BY fmp46Δ | *MATa his3Δ1 leu2Δ0 met15Δ0 ura3Δ0 fmp46Δ::kanMX4* | Open Biosystems |
| RKY219-1A | *MATa his3Δ1 leu2Δ0 lys2Δ0 met15Δ0 ura3Δ0 prx1Δ::kanMX4 fmp40Δ::kanMX4* | This study |
| RKY218-1B | *MATa his3Δ1 leu2Δ0 lys2Δ0 ura3Δ0 trx3Δ::kanMX4 fmp40Δ::kanMX4* | This study |
| RKY220-1C | *MATα his3Δ1 leu2Δ0 met15Δ0 ura3Δ0 grx2Δ::kanMX4 fmp40Δ::kanMX4* | This study |
| RKY344-1D | *MATa his3Δ1 leu2Δ0 met15Δ0 ura3Δ0 glr1Δ::kanMX4 fmp40Δ::kanMX4* | This study |
| RKY271-1C | *MATα his3Δ1 leu2Δ0 lys2Δ0 ura3Δ0 grx3Δ::kanMX4 fmp40Δ::kanMX4* | This study |
| RKY279-1C | *MATα his3Δ1 leu2Δ0 lys2Δ0 ura3Δ0 grx5Δ::kanMX4 fmp40Δ::kanMX4* | This study |
| RKY274-1A | *MATα his3Δ1 leu2Δ0 ura3Δ0 gpx1Δ::kanMX4 fmp40Δ::kanMX4* | This study |
| RKY345-2A | *MATa his3Δ1 leu2Δ0 met15Δ0 ura3Δ0 gpx2Δ::kanMX4 fmp40Δ::kanMX4* | This study |
| RKY346-1A | *MATa his3Δ1 leu2Δ0 met15Δ0 lys2Δ0 ura3Δ0 gpx3Δ::kanMX4 fmp40Δ::kanMX4* | This study |
| RKY217-1C | *MATα his3Δ1 leu2Δ0 ura3Δ0 lys2Δ0 gtt2Δ::kanMX4 fmp40Δ::kanMX4* | This study |
| RKY229-1A | *MATα his3Δ1 leu2Δ0 lys2Δ0 ura3Δ0 srx1Δ::kanMX4 fmp40Δ::kanMX4* | This study |
| RKY255-5A | *MATα his3Δ1 leu2Δ0 met15Δ0 lys2Δ0 ura3Δ0 gsh1Δ::kanMX4 fmp40Δ::kanMX4* | This study |
| RKY348-2B | *MATα his3Δ1 leu2Δ0 met15Δ0 lys2Δ0 ura3Δ0 yap1Δ::kanMX4 fmp40Δ::kanMX4* | This study |
| RKY276-1B | *MATa his3Δ1 leu2Δ0 lys2Δ0 ura3Δ0 cta1Δ::kanMX4 fmp40Δ::kanMX4* | This study |
| RKY275-1A | *MATα his3Δ1 leu2Δ0 lys2Δ0 ura3Δ0 ctt1Δ::kanMX4 fmp40Δ::kanMX4* | This study |
| RKY248-1D | *MATa his3Δ1 leu2Δ0 ura3Δ0 trr2Δ::kanMX4 fmp40Δ::kanMX4* | This study |
| RKY267-2A | *MATa his3Δ1 leu2Δ0 lys2Δ0 ura3Δ0 trx1Δ::kanMX4 fmp40Δ::kanMX4* | This study |
| RKY270-1C | *MATα his3Δ1 leu2Δ0 ura3Δ0 tsa1Δ::kanMX4 fmp40Δ::kanMX4* | This study |
| RKY230-1C | *MATα his3Δ1 leu2Δ0 lys2Δ0 ura3Δ0 tsa2Δ::kanMX4 fmp40Δ::kanMX4* | This study |
| RKY268-1B | *MATα his3Δ1 leu2Δ0 lys2Δ0 ura3Δ0 ahp1Δ::kanMX4 fmp40Δ::kanMX4* | This study |
| RKY278-1B | *MATa his3Δ1 leu2Δ0 met15Δ0 lys2Δ0 ura3Δ0 dot5Δ::kanMX4 fmp40Δ::kanMX4* | This study |
| RKY349-1A | *MATα his3Δ1 leu2Δ0 met15Δ0 lys2Δ0 ura3Δ0 sod1Δ::kanMX4 fmp40Δ::kanMX4* | This study |
| RKY350-3A | *MATα his3Δ1 leu2Δ0 met15Δ0 ura3Δ0 sod2Δ::kanMX4 fmp40Δ::kanMX4* | This study |
| RKY351 | *MATα his3Δ1 leu2Δ0 ura3Δ0 aim6Δ::kanMX4 fmp40Δ::kanMX4* | This study |
| RKY352 | *MATα his3Δ1 leu2Δ0 lys2Δ0 ura3Δ0 aim14Δ::kanMX4 fmp40Δ::kanMX4* | This study |
| RKY269-1C | *MATa his3Δ1 leu2Δ0 met15Δ0 ura3Δ0 aim33Δ::kanMX4 fmp40Δ::kanMX4* | This study |
| RKY272-3A | *MATα his3Δ1 leu2Δ0 met15Δ0 ura3Δ0 oye2Δ::kanMX4 fmp40Δ::kanMX4* | This study |
| RKY273-1B | *MATα his3Δ1 leu2Δ0 met15Δ0 ura3Δ0 ycp4Δ::kanMX4 fmp40Δ::kanMX4* | This study |
| RKY353-2A | *MATa his3Δ1 leu2Δ0 met15Δ0 lys2Δ0 ura3Δ0 mdl2Δ::kanMX4 fmp40Δ::kanMX4* | This study |
| RKY266-1 | *MATα his3Δ1 leu2Δ0 ura3Δ0 aif1Δ::kanMX4 fmp40Δ::kanMX4* | This study |
| RKY277-1B | *MATa his3Δ1 leu2Δ0 met15Δ0 ura3Δ0 mgp12Δ::kanMX4 fmp40Δ::kanMX4* | This study |
| RKY354-3A | *MATα his3Δ1 leu2Δ0 lys2Δ0 ura3Δ0 oxr1Δ::kanMX4 fmp40Δ::kanMX4* | This study |
| RKY355-1C | *MATa his3Δ1 leu2Δ0 lys2Δ0 ura3Δ0 pos5Δ::kanMX4 fmp40Δ::kanMX4* | This study |
| CPY17-1 | *MATα his3Δ1 leu2Δ0 met15Δ0 lys2Δ0 ura3Δ0 fmp46Δ::kanMX4 fmp40Δ::kanMX4* | This study |
| RKY357 | *MATa his3Δ1 leu2Δ0 met15Δ0 ura3Δ0 TRX3-T66>E* | This study |
| RKY360 | *MATa his3Δ1 leu2Δ0 met15Δ0 ura3Δ0 TRX3-T66>A* | This study |

### **Table S2**. Primers used in the study.

| Name | sequence (5'→3') |
|---|---|
| Primers for gene replacement cassettes amplification, tagging, mutagenesis and for clone verification by PCR | |
| Prx1-Up | GATATTCCTGATAGCCACTATGT |
| Prx1-Low | TGACATTAAGGTTTCTGCAGAGAG |
| Prx1-Ver | TTGACCTCGAGCAAGCGCTCCACTAT |
| Trx3-Up | GAAGAACAACGCCCTAGATATAGTGA |
| Trx3-Low | CAGTTACTTTCCTCACCATTCAT |
| Trx3-Ver | GTATGAAGATGGCTTATTATGATC |
| Trx3-For | GCTAGCCCCCTAACCAAAGAAAGC |
| Trx3-T66E | CCCTGTAAGATGATGCAACCACACTTA**GAG**AA**G**TTAATTCAGGCTTATCCAGATGTAAG |
| Trx3-T66A | CCCTGTAAGATGATGCAACCACACTTA**GCG**AA**G**TTAATTCAGGCTTATCCAGATGTAAG |
| Trx3-Mut-left | GCCACACCAAGTAGCATAAAAGTCG |

| | |
|---|---|
| Trx3-3HA-Up | GGCCAACTCATCGGCAAGATCATTGGAGCTAACCCTACTGCTTTAGAGAAGGGAATCAAA GATCTACGGATCCCCGGGTTAATTAA |
| Trx3-3HA-Low | CTATCTTTTTTCTTATATTTAAAACTAAAAAAAAAAAAACTACATGTGCTCATGAATATAAAA TGAATTCGAGCTCGTTTA |
| Grx2-Up | CAGAACTATTACTAAGATCCG |
| Grx2-Low | GGATCTTGTACCTCAGACGG |
| Grx2-Ver | GCCTCCCTCTTGATTTATGC |
| Yca1-Low | GTCAAGATCACCGTTGAGC |
| Yca1-Low | CGCTCGTTCCACACCTAATAG |
| Yca1-Ver | AAGAGCATCTTTGAACTCTG |
| Fmp40-Up | CGGTGATATGAGGTGATCGTGG |
| Fmp40-Low | GGTGCCAGTCGTTCGGCTACCC |
| Fmp40-Ver | GAACTCCGGAATTGGACGATG |
| Verif-KanMx | GGATGTATGGGCTAAATGTACG |
| Ctt1lower | AATAATTATGGAGATATAATTACGAATAATTATGAATAAATAGTGCTGCCGAATTCGAGCTC GTTTAAAC |
| Ctt1upper | CTGTTGACCTTGAAGGTTATGCCAAGACTTGGTCCATTGCAAGTGCCAATCGGATCCCCG GGTTAATTAAC |
| Cta1lower | CATAAATAATTGTCGTGGAAACAACGCCACTCATTTGTTACTTGAGCGTTGAATTCGAGCT CGTTTAAAC |
| Cta1upper | AAGTAGCTGAGGCAAAACATGCTTCTGAGCTTTCGAGTAACTCCAAATTTCGGATCCCCG GGTTAATTAAC |
| Prx1-S116A-Up | AGAAATGTTAAATTGATCGGGCTTGCTGTGGAAGATGTTGAGTCCCACG |
| Prx1-S116A-Low | CTTGTCGAATTCCGGCTTC |
| Trx3-T95A-Up | CAAAGAGTGTGAAGTGGCGGCTATGCCCACCTTTGTTCTT |
| Trx3-T95A-Low | GCAATATCTGGTGATTCG |
| Trr2-Ver | CAACCGACTATTCTTCTCATCG |
| Glr1-Ver | GACGTCACTTAGTAAGCACG |
| Gpx2-Ver | GCAAAGCCCCTAAGCCAGCC |
| Cta1-Ver | AAGTTTGTGAAGAAACCATCG |
| Ctt1-Ver | TACGTCGCCGATCCAGAAG |
| Ycp4-Ver | CCTTGGCTAGCGTACTTATTTCG |
| Yap1-Ver | AACTGATGGATGCGTAAGGTC |
| Aif1-Ver | TGTCCACCCTGCATGATTAGTC |
| Gpx3-Ver | CCATGAGGAACAGTATGCTC |
| Aim6-Ver | CCTCTTCAAAGGACAGCCC |
| Aim14-Ver | GGTATAATACACAGCCGGG |
| Mdl2-Ver | CGGGAAACTAATAATGACTAC |
| Oxr1-Ver | CAATGGCAGTATGAAAGCCTG |
| Gsh2-Ver | GTGTCATTTCCATCAGTACGC |
| Pos5-Ver | GCTGCATGTATCTATATGCATC |
| Trx1-Ver | CCGATGAATTAGTGGAACCAAG |
| Ahp1-Ver | CAATTTCCTCTCCCTTGCC |
| Aim33-Ver | TAGATCCATCTACAGGGAAC |
| Tsa1-Ver | TGGCTCGGGTTGGCAAAGTC |
| Grx3-Ver | GAGAATGTGTGCCGGGTCATG |
| Oye2-Ver | GCCTTGTTCCTCCTGGTGAC |
| Gpx1-Ver | GGCTCACCTCTACTACAGAG |
| Mgp12-Ver | GGGTCTATGATGTGCGCAAG |
| Dot5-Ver | GAATGTTCGAGCAAGAGCATG |
| Grx5-Ver | GTGCCGTAAAGAGTCTAAACCGG |
| Trr1-Ver | CCACGCGTATAAGAGTGGAC |
| Gtt2-Ver | TAGGCGCTGGAAATGACCCG |
| Tsa2-Ver | CAAGGATCAACTATGGATGAG |
| Srx1-Ver | AGGCCGGCCCAGAAGTCGCC |
| Fmp46-Ver | GTGATGTACCATGGCCTATTCA |
| Sod1-Ver | GAAGTACAGAACCCGCTCCC |
| Sod2-Ver | CTAATTGCTATTATCATTGTTGGCG |
| Gsh1-Ver | TGAAGCTTGTTCTTGCCTC |
| **Primers for qPCR analysis** | |
| ScAHP1cF | GGCTGTGCTTTCACCAAATC |
| ScAHP1cR | TCCTTGGCAGCGTAAGTAAC |
| ScAIF1aF | GGACTCTGGGTTATACAACGATAC |
| ScAIF1aR | TTTCGGCGAGGTGTCTAAAG |

| | |
|---|---|
| ScCCP1aF | TCTTGCTTGGCACACTTCAG |
| ScCCP1aR | TTGAAGCCATTCTGCAAGCC |
| ScCTA1bF | CCCTTATCATTGGGCAACATCC |
| ScCTA1bR | TGCCGATGTTATATGCCAAGTTC |
| ScCTT1bF | CACCCTAACGGAGAATGTTGAC |
| ScCTT1bR | GTCTGGCTTGTAGAACGGAATC |
| ScDOT5cF | ATGCTGCTGTCTTTGGACTG |
| ScDOT5cR | CCCAATAAACTCTCTCTTTGGATCG |
| ScFM46bF | CCACGTACCATCTCGTTGTTTAC |
| ScFM46bR | AAGCGGTTGGCGATTTCC |
| ScGLR1bF | GTATTGAGCTAGCAGGTGTGTTC |
| ScGLR1bR | GGTCAGTAATAGTGTTCTGGATGC |
| ScGPX1bF | TCCCTGCGTAACAAAGTGG |
| ScGPX1bR | AAAGGCCACAATCACTAGACC |
| ScGPX2aF | TTCACGCCGCAGTATAAAG |
| ScGPX2aR | CTGCTTCCCGAACTGATTAC |
| ScGPX3aF | GGATTTACCATCATCGGGTTC |
| ScGPX3aR | GGAAAGTCACGCCATAGTTC |
| ScGRX1bF | GTACTGTCCATACTGCCATGC |
| ScGRX1bR | AATGTCTGCGCCTTCCTTC |
| ScGRX2bF | AAAGACATACTGCCCTTACTG |
| ScGRX2bR | GAGCCATTGCTCATTTCATC |
| ScGRX3bF | CTCAGTCAATTCCGGATCATCAC |
| ScGRX3bR | CCTCAGTTTCTTCTTCCTCTTCTTC |
| ScGRX5cF | TTCAGAATGGCCAACTATTCCAC |
| ScGRX5cR | CCTGTGCCTCTTCTAGCAAATC |
| ScGRX6aF | AGGGTCGCGAATAACCAAAG |
| ScGRX6aR | TCAAGCAGTTCCTTCATGCC |
| ScGRX8cF | TCACTTCCAAGAAACGAACAGG |
| ScGRX8cR | CTATGCAATTGACTCTCAGTACCC |
| ScGSH1aF | CGTGCCATTGCAGAATATAAG |
| ScGSH1aR | AAGACAGTCAGCGTCAAAG |
| ScGSH2bF | ACTGGGAGGCAAGACTATTCC |
| ScGSH2bR | ACCTAATACGCCCTCATCTGTC |
| ScGTT1cF | AGTCCATTGGTTGGACCATTC |
| ScGTT1cR | GAGCACGGAAGTTAGCATCTC |
| ScGTT2aF | AGTGCCAGTGCTTGAACTTG |
| ScGTT2aR | TTTTCCAGCGGTGTTTTGCC |
| ScMSN2aF | ACGGTGCCGCTTACAAATC |
| ScMSN2aR | CCTGTAACTTGGCCTTCTTTCC |
| ScMSN4aF | CCATAGCAGTAGCACCACAAG |
| ScMSN4aR | CAGACAGAGGATTTGCCATCAC |
| ScOXR1aF | TTTATGCGATGACGGCCTAC |
| ScOXR1aR | ACGCCATACTTCCAAAGCTAC |
| ScPOS5aF | CATAGATGGTACTCAGCTTCCG |
| ScPOS5aR | GGTCTTGGCGACACAATAAATC |
| ScPRX1bF | TTTACGACTACTTGGGCGACTC |
| ScPRX1bR | TCGAATTCCGGCTTCAATTTGG |
| ScPST2bF | GGTGCTCCAAAGCCAGATTAC |
| ScPST2bR | GTCCCAGAAAGCCTTCCATTG |
| ScSKN7aF | GCCATTGCTAATGCTAGGTCAG |
| ScSKN7aR | ATTGTGGTAATTGCGGAGTGG |
| ScSOD1aF | TTCGAACAGGCTTCCGAATC |
| ScSOD1aR | AGCAGAGACACAACCATTGG |
| ScSOD2cF | GTTCCTCTAGTTGCCATTGACG |
| ScSOD2cR | TGGATGCTTCTTTCCAGTTGAC |
| ScSRX1bF | ACCAACAGAAATTCCTCTCTCAG |
| ScSRX1bR | TTGCTAGCGGTGGGAATG |
| ScTDP1aF | CTGGGAGGAAGAAAGCAGAAATAC |
| ScTDP1aR | CTTTAAGGGACTGGTGACAAACTC |
| ScTRR1bF | ATCCACGAAGTTTGGCACTG |
| ScTRR1bR | GTCACAGGTTCTGCGTCTTC |
| ScTRR2cF | AGAGCCTGTGACCACTGATG |
| ScTRR2cR | CACATACAGCACAGGCAGATATTC |

| | |
|---|---|
| ScTRX1cF | GGATGTCGATGAATTGGGTGATG |
| ScTRX1cR | GCTTAATAGCCGCTGGGTTG |
| ScTRX3bF | CCAGATATTGCCAAAGAGTGTGAAG |
| ScTRX3bR | CCCTTCTCTAAAGCAGTAGGGTTAG |
| ScTSA1bF | CCCAAGAAAGGAAGGTGGTTTG |
| ScTSA1bR | CAAACCTCTCAAGGCGACAC |
| ScTSA2bF | CGGTGGATTAGGTCCAGTTAAAG |
| ScTSA2bR | CGGGTCGATTATGAACAAACCTC |
| ScYAP1aF | AATGCTGCCTCTAACACCAATC |
| ScYAP1aR | AGTTGAATTGTCGTCTGGGAAAG |
| ScYCP4cF | GGTCGAGGAAACTTTACCTGATG |
| ScYCP4cR | AAGGCGTCATATTCGAGCAAC |
| ScYHB1bF | CGGTAAACACTCACCCAATTAGAC |
| ScYHB1bR | CTCTGGCAGCTTCCATCTTTAC |
| ScLSR1bF | CGACGGTTGCTCAAGGTTATTG |
| ScLSR1bR | TCGGAAAGACAGGGAAGAGTATG |

### Supplementary Figures

A

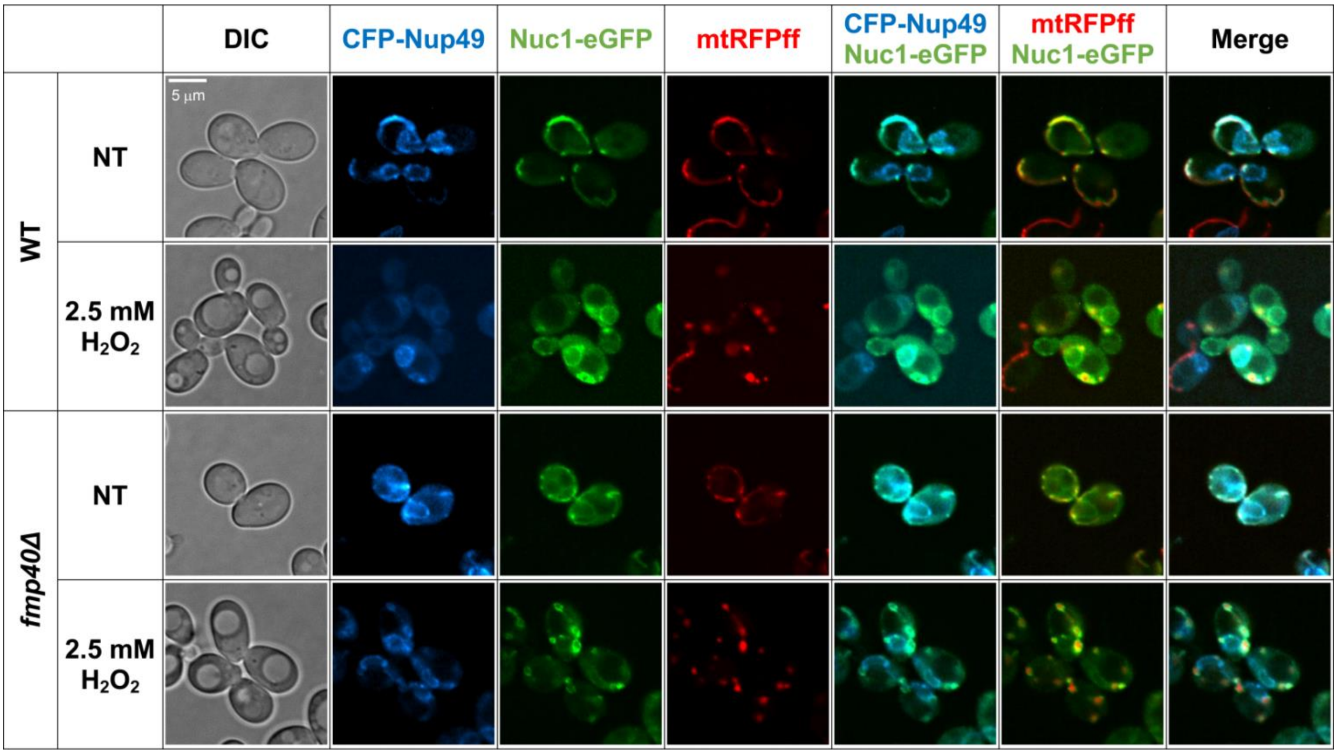


B

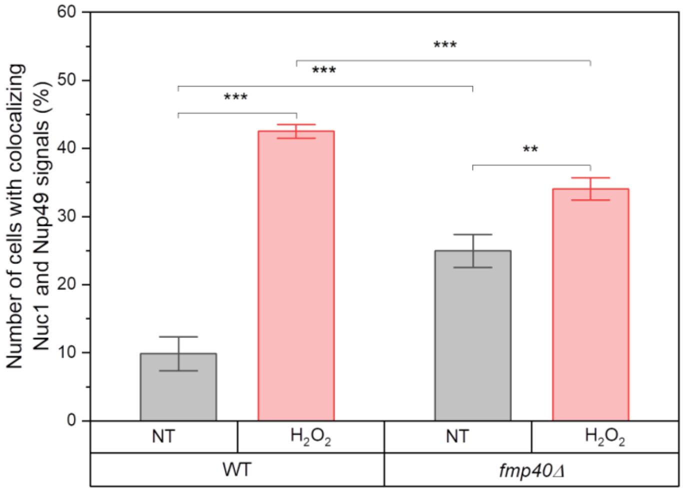


**Fig. S1.** Nuc1 localizes predominantly in the mitochondria in the control conditions, but after hydrogen peroxide treatment, it efficiently translocates to the nucleus; however, in the *fmp40Δ* cells, this translocation occurs less frequently. (**A**) The strains carrying plasmids: pWJ1323 [*CFP::NUP49*] [3] and pNuc1-GFP [*TEF1pNUC1::eGFP*] and pYX142-mt-RFPff [4] were cultivated to exponential growth phase in minimal selection medium (SC-ura-his-leu) with shaking then treated with 2.5 mM $H_2O_2$ for 200 minutes to induce PCD. The cells' images were collected using fluorescence microscopy. Representative examples of cellular localization of Nuc1-eGFP, nucleus marker CFP-Nup49, and mitochondrial marker mt-RFPff are shown for all tested strains and conditions. (**B**) The frequency of Nuc1 and Nup49 co-localization was quantified based on fluorescent microscopy studies. Approximately 600 cells (~200 per each of three biological repetition) containing all fluorescent signals, per each strain and conditions were analyzed. To check if the results were statistically significant, the t-Welch test was applied using Origin Pro software. P-values (** - p-value < 0.01; *** - p-value < 0.001) indicate that there is a statistically significant association between variables.

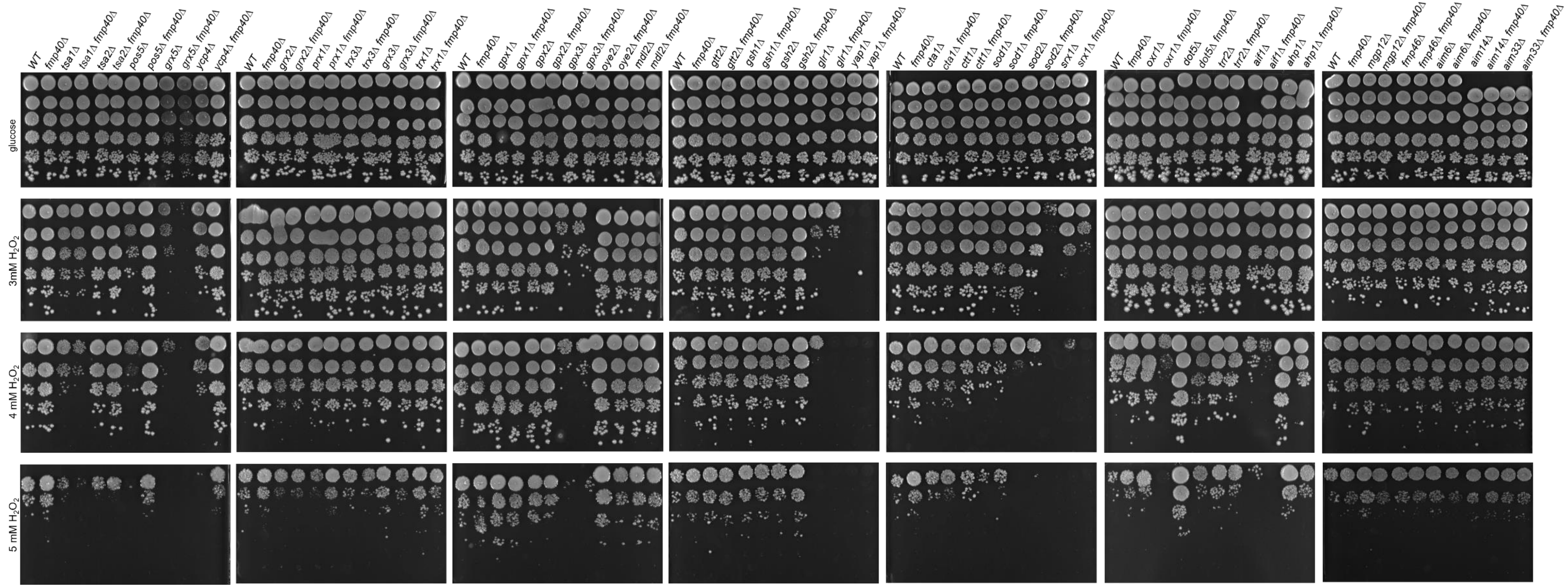


**Fig. S2. The genetic interactions of the *fmp40Δ* cells with the indicated genes encoding known redox proteins, assessed by the growth phenotype in the presence of hydrogen peroxide**. Cells from the indicated strains grown in YPGlu medium were serially diluted and spotted on YPGlu solid medium without or with given concentration of hydrogen peroxide and incubated at 28°C. The plates were photographed after three days of incubation. Experiment was performed at triplicate and representative plates are shown.

...

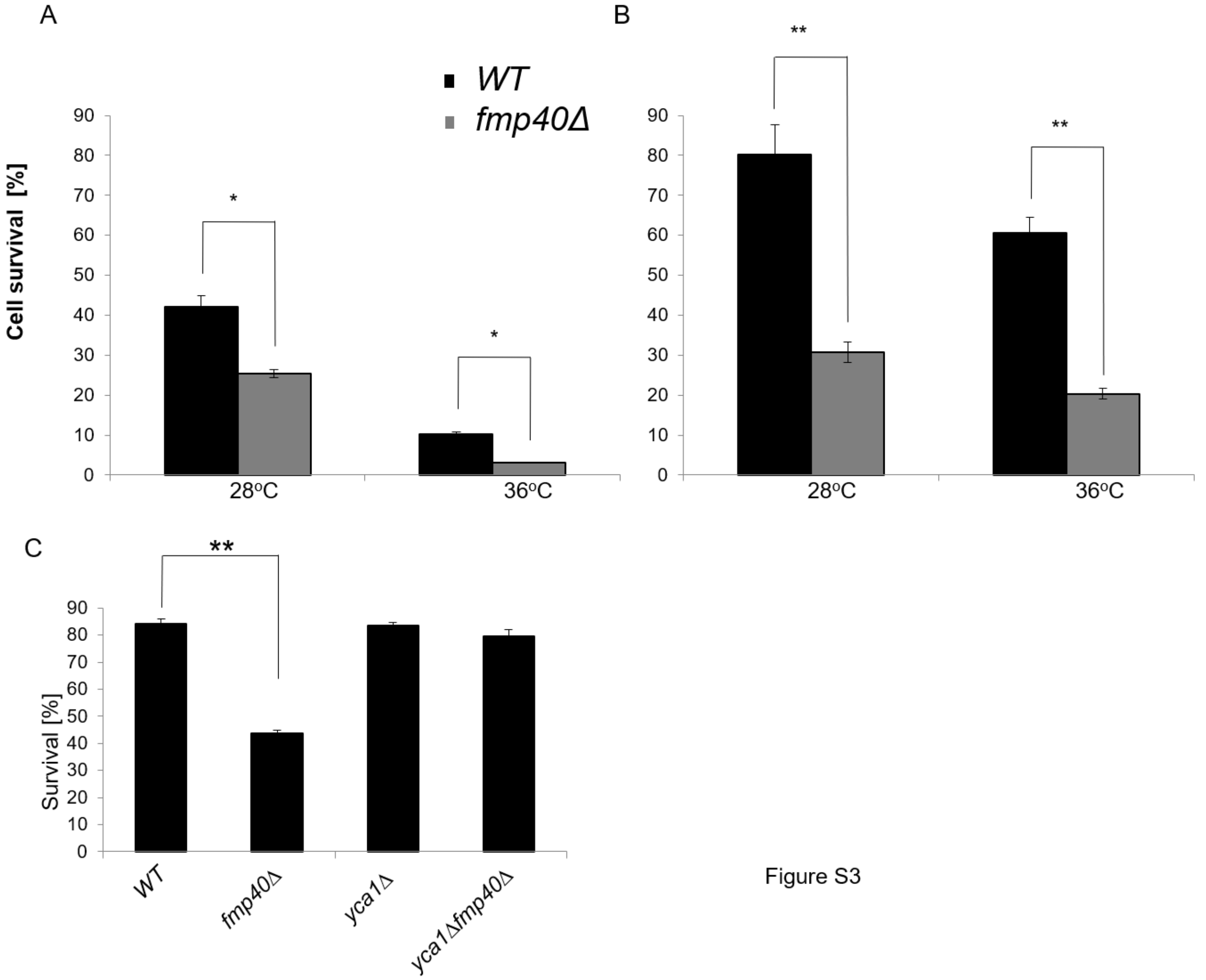


**Fig. S3. The decreased survival of *fmp40Δ* cells in the presence of hydrogenof hydrogen peroxide in different media.** A liquid rich complete medium containing glucose **(A)** or glycerol **(B,C)** as a carbon source was inoculated with the indicated strains in W303-1B background and grown over night with shaking at 28 °C until the optical density $OD_{600}$=3. Then the cultures were divided into two of equal volume and to one the hydrogen peroxide was added to 0.2 mM concentration for the glucose medium or 0.03 mM for the glycerol medium. Both cultures were incubated for 200 min with shaking at 36°C or 28°C. Cell viability was then determined according to the colony forming unit method taking the amount of growing colonies in non-treated culture as 100%.

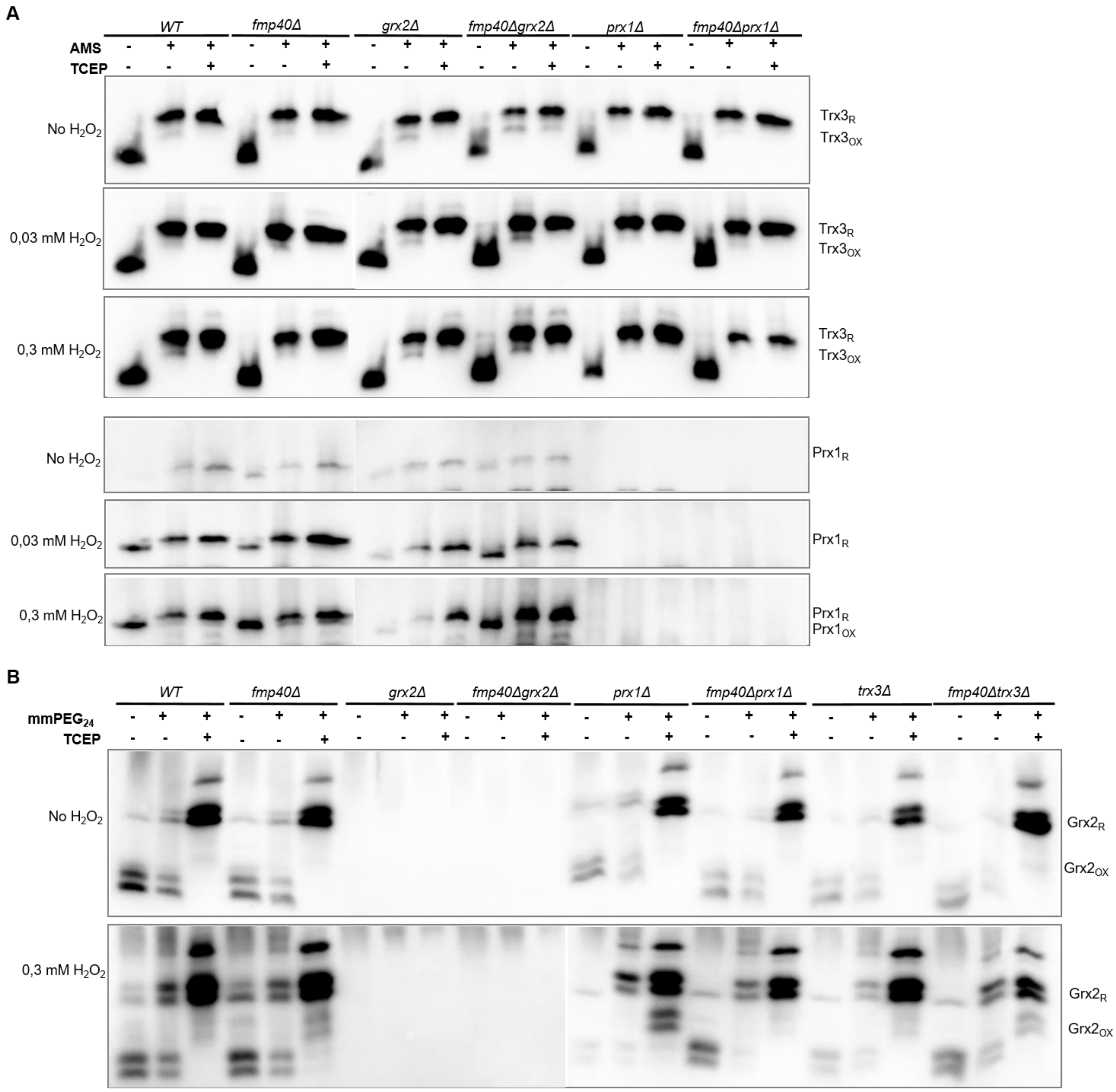


**Fig. S4. Redox shift assays of Trx3 and Prx1 proteins. (A)** In the indicated strains *TRX3::Myc* gene fusion was expressed from the natural *TRX3* genomic locus. Strains were grown to the exponential phase in YPGlyA medium and subsequently treated with the indicated concentrations of $H_2O_2$ for 200 min. Proteins were extracted, incubated with AMS or TCEP when indicated, separated in 18% SDS-PAGE gels, transferred to the membrane and immunodetected with anti-Myc or anti-Prx1 antibodies. **(B)** The indicated strains were grown as above, proteins were extracted, incubated with TCEP or $mmPEG_{24}$. Proteins were then analyzed in 16% SDS-PAGE and after transfer to the membrane immunodetected using anti-Grx2 antibody.

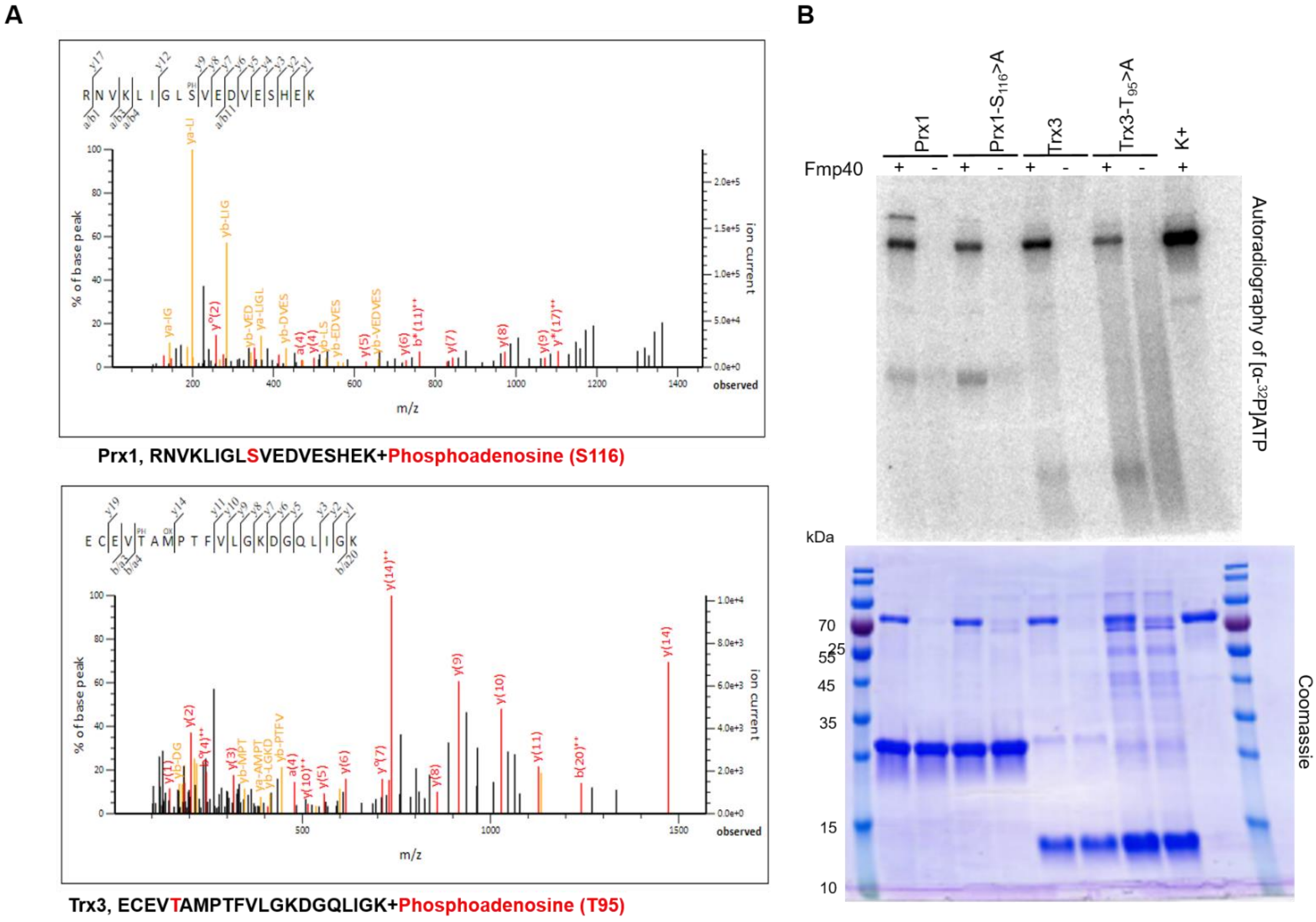


**Fig. S5. AMPylated residues identified in Prx1 and Trx3 proteins in *in vitro* AMPylation reactions with Fmp40** (related to Fig. 8). **(A)** MS/MS spectra of Prx1 and Trx3 peptides ions detected in the reactions performed with Fmp40. The precursor ions, m/z 794.4 (3+) and m/z 876.4 (3+), were subjected to HCD fragmentation to generate the MS/MS spectra shown. Unique ion corresponding to neutral loss of the AMP group is present at 136.1. Peaks labeled with a single asterisk (*) correspond to neutral loss of ammonia (−17 Da) from fragment ions. **(B)** Autoradiograph depicting the incorporation of $\alpha{-}^{32}P$ from $[\alpha{-}^{32}P]$ATP into the Prx1 and Trx3 recombinant proteins variants, purified from *E. coli* cells, by Fmp40 (see Materials and Methods for experimental details). Auto-AMPylation of Fmp40 was used as a positive control. Reactions products were separated by SDS-PAGE and visualized by Coomassie blue staining (lower panel) and autoradiography (upper panel).

```
1     MLFYKPVMRM  AVRPLKSIRF  QSSYTSITKL  TNLTEFRNLI  KQNDKLVIDF  YATWCGPCKM
61    MQPHLTKLIQ  AYPDVRFVKC  DVDESPDIAK  ECEVTAMPTF  VLGKDGQLIG  KIIGANPTAL
121   EKGIKDL*
```

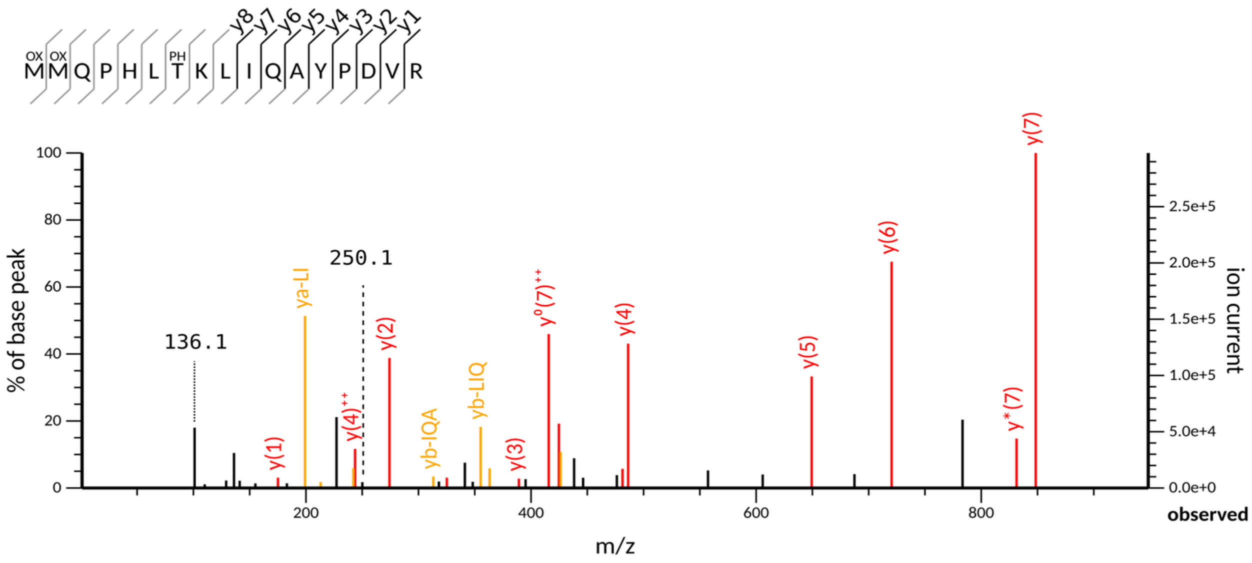


**Fig. S6. AMPylated residue identified in Trx3 protein purified by affinity chromatography from freshly isolated native mitochondria.** The precursor ion m/z 801.37 (3+) and m/z 499.71 (2+) was subjected to HCD fragmentation to generate the MS/MS spectrum shown. Unique ions corresponding to neutral loss of the AMP group are present at 136.1 and 250.1 Da. Unique ions corresponding to neutral loss of the AMP group (labeled with **) are also present at 136.1 and 250.1 Da. Peaks labeled with a single asterisk (*) in both spectra correspond to neutral loss of ammonia (−17 Da) or water (−18 Da) from fragment ions.

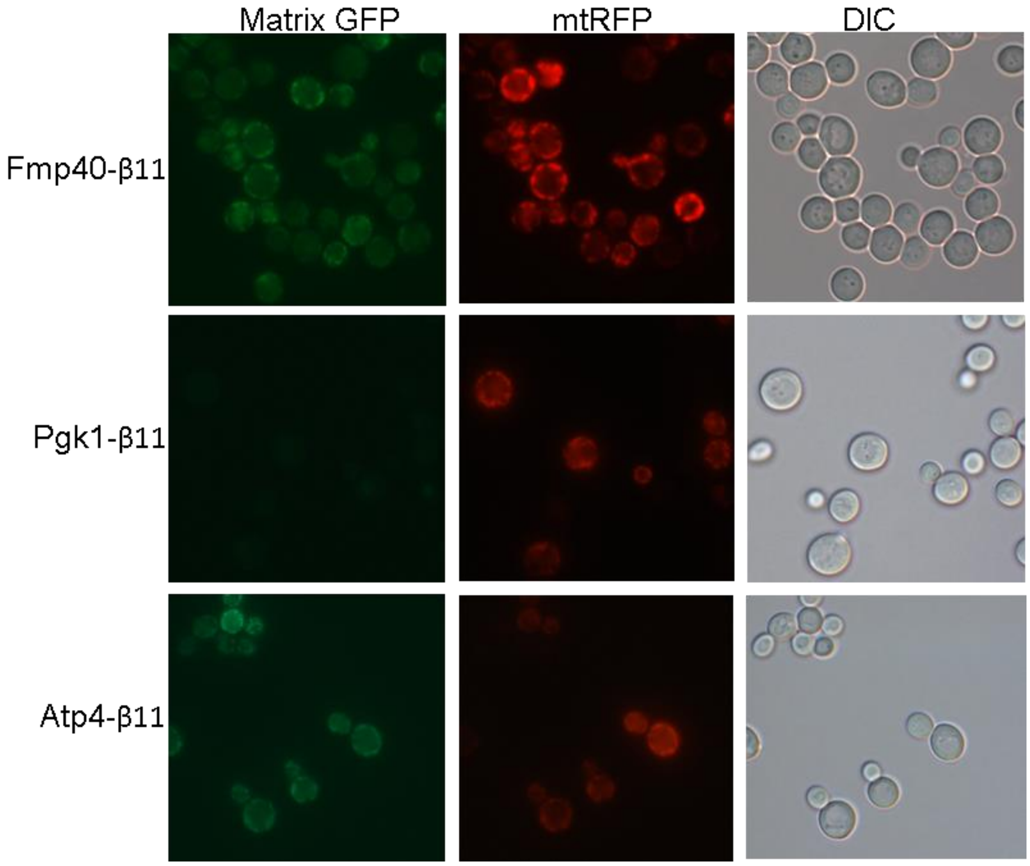


**Fig. S7. Fmp40 have the matrix echoform.** Fluorescence microscopy analyzes of BiG Mito-Split-GFP strain transformed with pAG414pGPDβ11 expressing Fmp40-β11, Pgk1-β11 or Atp4-β11 [2]. Cells were grown in SC+glicerol-ura liquid medium till $OD_{600}$=2 and GFP reconstitution upon mitochondrial import was followed by epifluorescence microscopy (N = 3).

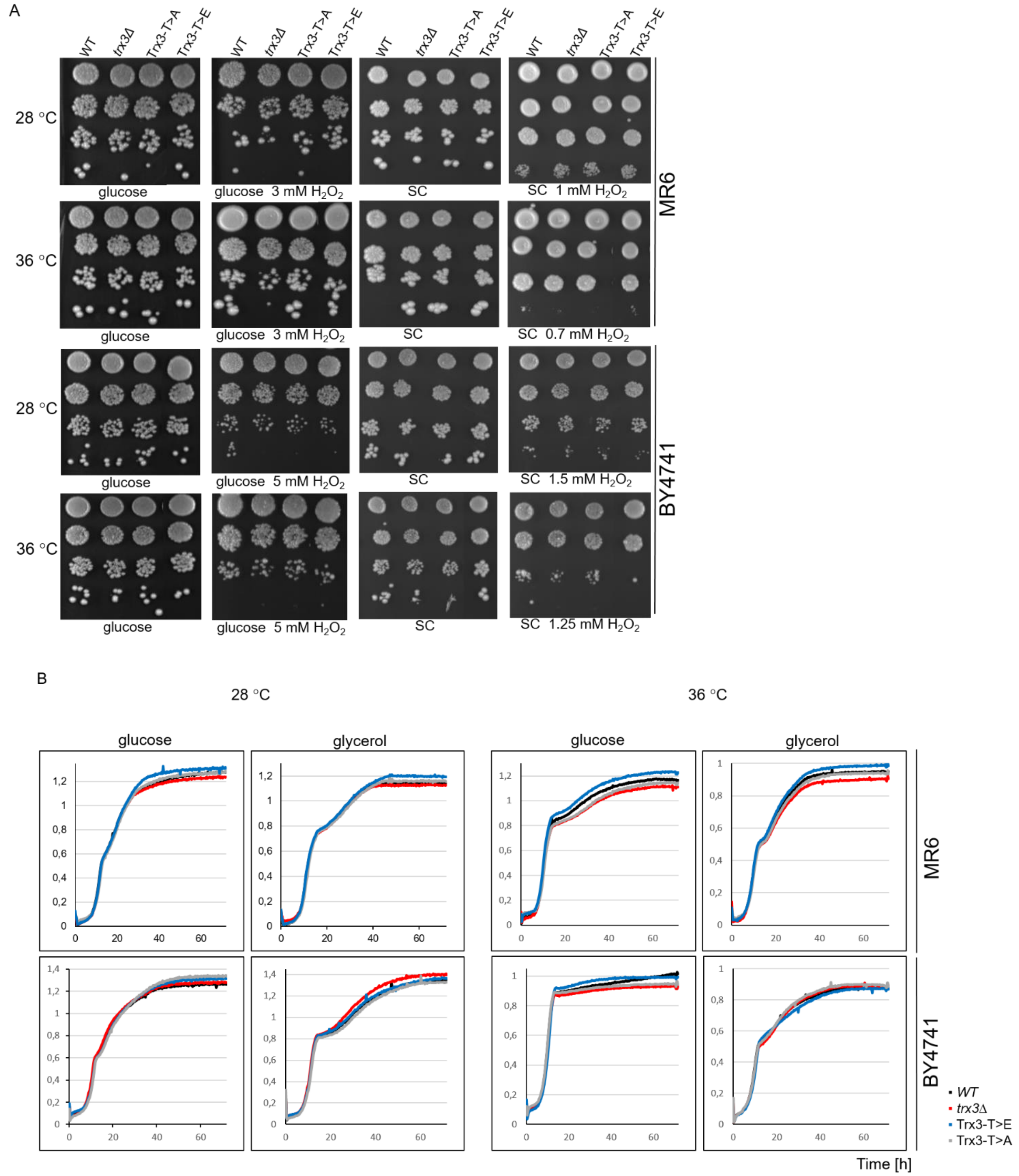


**Fig. S8. Influence of amino acid substitutions in Thr66 of Trx3 on the growth phenotype of yeast carrying respective variants**. The growth of yeast cells expressing the *TRX3* variants in fermentative and respiratory solid (**A**) or liquid (**B**) media. Cells from the indicated strains grown in liquid YPGlu medium were serially diluted and spotted on YPGluA or YPGlyA containing plates (A) or inoculated at $OD_{600}$=0.1 and grown using a Bioscreen CTM system at 28 or 36 °C, with shaking. The plates were photographed after three days of incubation in respective temperature. Experiments was performed in triplicate and representative photographs and curves are shown.

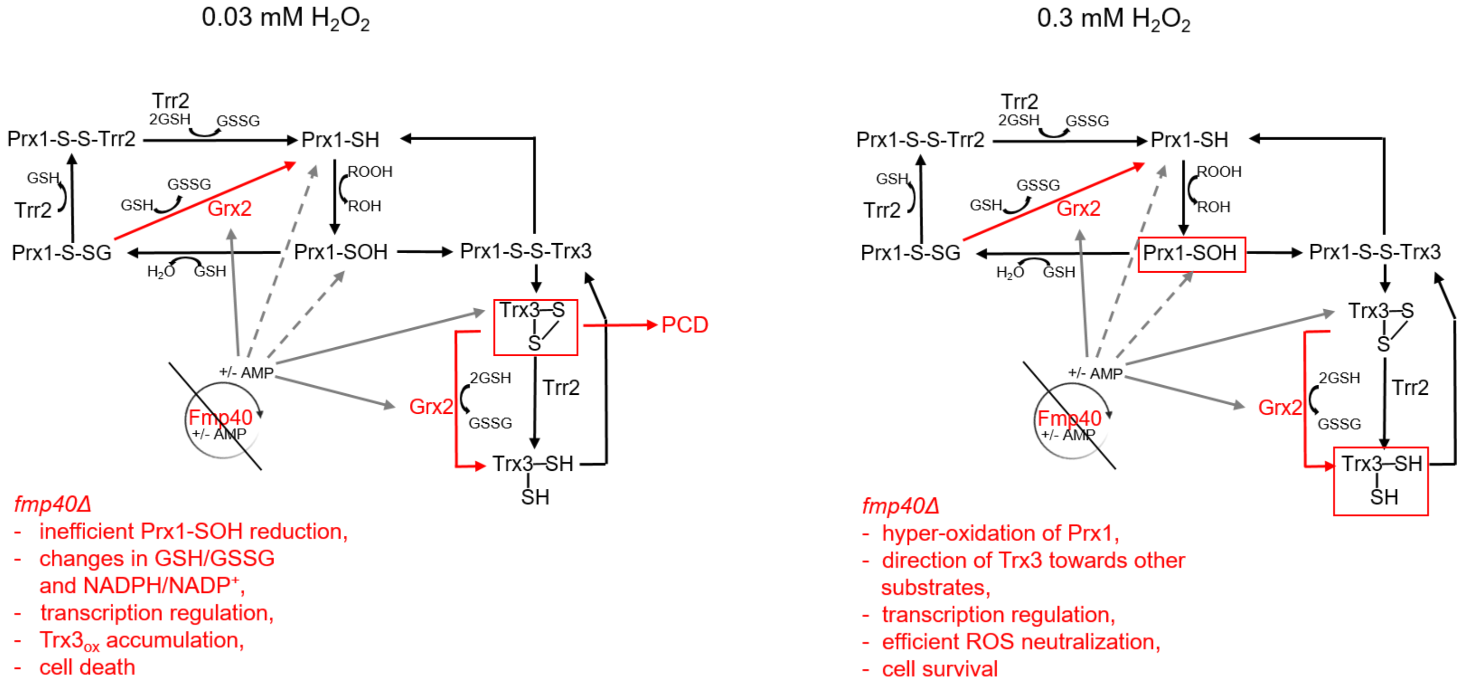


**Fig. S9. Mechanisms of PCD inhibition/induction in *fmp40Δ* cells in response to oxidative stress.** In the absence of Fmp40, upon treatment with low hydrogen peroxide concentrations, due to the ineffective reduction of Trx3, the balance of GSH/GSSG and NADPH/NADP$^+$ is changed, the expression of many genes encoding proteins necessary for ROS neutralization/signaling decreases, the Trx3-S-S accumulates, and PCD is triggered. At high concentrations of hydrogen peroxide, in *fmp40Δ* cells, the Prx1 pool is rapidly hyper-oxidized, so it neither degrades hydrogen peroxide nor signals the oxidation of ROS effectors. Hyperoxidation of Prx1 leads to the targeting of reduced Trx3 to other substrates, influencing the adaptive transcriptional response and enabling the reduction of oxidized proteins, which is necessary for cell survival. In parallel, the pool of GSH decreases without changes in the level of GSSG, which indicates its consumption for the reduction processes other than Prx1 reduction. This results in the effective removal of ROS and in higher viability.
